# Rupture Strength of Living Cell Monolayers

**DOI:** 10.1101/2023.01.05.522736

**Authors:** Julia Duque, Alessandra Bonfanti, Jonathan Fouchard, Lucia Baldauf, Sara R. Azenha, Emma Ferber, Andrew Harris, Elias H. Barriga, Alexandre J. Kabla, Guillaume Charras

## Abstract

The ability of tissues to sustain mechanical stress and avoid rupture is a fundamental pillar of their function. Rupture in response to physiological levels of stress can be undesired, for example resulting from disease or genetic mutations, or be an integral part of developmental processes, such as during blastocoel formation in mouse or leg eversion in flies. Despite its importance, we know very little about rupture in cellularised tissues because it is a multi-scale phenomenon that necessitates comprehension of the interplay between mechanical forces and processes at the molecular and cellular scales. Using a combination of mechanical measurements, live imaging and computational modelling, we characterise rupture in epithelial monolayers. We show that, despite consisting of only a single layer of cells, monolayers can withstand surprisingly large deformations, often accommodating several-fold increases in their length before rupture. At large deformation, epithelia increase their stiffness multiple-fold in a process controlled by a supracellular network of keratin filaments. Perturbing keratin organisation fragilised monolayers and prevented strain stiffening. Using computational approaches, we show that, although the kinetics of adhesive bond rupture ultimately control tissue strength, tissue rheology and the history of deformation prior to failure set the strain and stress that the tissue reaches at the onset of fracture. Our data paint a picture of epithelia as versatile materials that combine resistance to shocks with deformability when subjected to low strain rates.

## Introduction

During development and normal physiological function in adult tissues, epithelial monolayers withstand mechanical stresses that differ markedly. In embryonic morphogenesis, tissues undergo large deformations over periods of hours or days. For example, convergence and extension in *Xenopus Laevis* typically leads to a ∼3 fold increase in tissue length over a span of ten hours, a process enabled by rearrangement of adhesive contacts at the cellular-scale. In contrast, deformations in adult tissues are typically smaller and take place on shorter time scales with fixed tissue organisation. For example, the myocardium tissue deforms by 10 - 20% in 200 ms [1], lung alveoli deform by ∼ 20% up 20 times a minute during breathing [2], and the skin deforms by over 50% in fractions of a second during limb movement [3, 4]. The mechanical role of epithelia is particularly apparent in disease. Mutations in intermediate filaments and desmosomal proteins in the epidermis lead to *epidermolysis bullosa*, a family of diseases characterised by fragile skin that fractures in response to physiological levels of deformation [5, 6]. Despite its importance, we know relatively little about the strength of epithelia and the biological structures that control it, partly because of the difficulty of characterising fracture in living cellularised tissues.

Fracture is a permanent break of a material into smaller components when subjected to stress. Loss of material integrity can occur once a threshold of strain or stress is exceeded and, the mode of fracture often depends on the rate of stress application. While much is known about fracture in classic engineering materials such as ceramics and steel [7, 8], fracture in soft materials is still an active research field [9, 10, 11]. Tissue fracture is inherently a multi-scale process with deformations applied at the tissue-scale resulting in stress at the cellular-scale that causes rupture of intercellular adhesion complexes at the molecular-scale. Further complexity arises because of the viscoelastic properties of living tissues that stem from biological processes with distinct time-scales, such as protein turnover occurring over tens of seconds to tens of minutes, cell intercalation over tens of minutes to hours, and upregulation of cell division taking hours. Thus, fracture in living tissues involves processes spanning multiple length- and time-scales.

Rupture can occur either in response to extrinsically applied forces or because of forces generated by the cells within a tissue. While we are familiar with the former from our everyday experience, the latter is unique to living tissues and is a normal part of some developmental processes. Indeed, tissues can self-rupture as a consequence of local upregulation of active stress due to myosin contractility, motility on a substrate, or an increase in osmotic forces to ensure the proper development of embryonic structures. For example, during eversion of the *Drosophila* leg, the squamous peripodial epithelium first delaminates from its collagenous substrate before rupturing in response to a local increase in cellular contractility [12]. *Trichoplax adhaerens*, a flat marine metazoan, reproduces asexually by fission induced by movement of different parts of its body in opposite directions [13]. During blastocoel formation in the mouse, cracks occur between the inner mass cells in response to osmotic pumping [14, 15]. In gastrulation, tissue integrity is transiently lost because of an epithelial to mesenchymal transition (EMT) that is thought to involve an upregulation in cell contractility [16]. Excess intrinsic stresses can also result in unwanted rupture. For example, depletion of myosin phosphatase leads to excessive contractility in the *Drosophila* amnioserosa, giving rise to frequent breaks between cells [17]. At the onset of metastasis, tumour cells undergo EMT in a process dependent on myosin contractility [18]. Collectively, these phenomena suggest that tissues have the intrinsic ability to self-rupture. However, it remains unclear if ruptures arising from extrinsic and intrinsic forces occur through the same biophysical processes.

Here, by using suspended epithelial monolayers of MDCK cells, we investigate rupture at the tissue-scale by subjecting tissues to ramps in deformation. We compare ruptures occurring in response to increases in contractility to those due to externally applied deformation. At large deformation, we reveal that epithelia strain-stiffen and, surprisingly, actomyosin is not the main contributor to mechanics, rather this is dominated by a supracellular network of keratin filaments. Finally, using computational modelling, we link strain at the tissue level to rupture of intercellular adhesion bonds at the molecular scale to predict the onset of tissue fracture. We show that tissue-scale rheology interplays with molecular bond rupture kinetics to govern tissue rupture.

## Results

### Epithelial monolayers can withstand large deformations before rupture

To investigate the response of living tissues to externally applied deformation, we used Madin-Darby canine kidney (MDCK) monolayers devoid of a substrate and suspended between two test rods [19]. In these conditions, all of the force applied to the monolayer is borne by intercellular adhesions and transmitted through the cytoskeleton (**Fig. 1a**), making this an ideal system to explore the operating limits of a living cellularised material.

**Figure 1:**
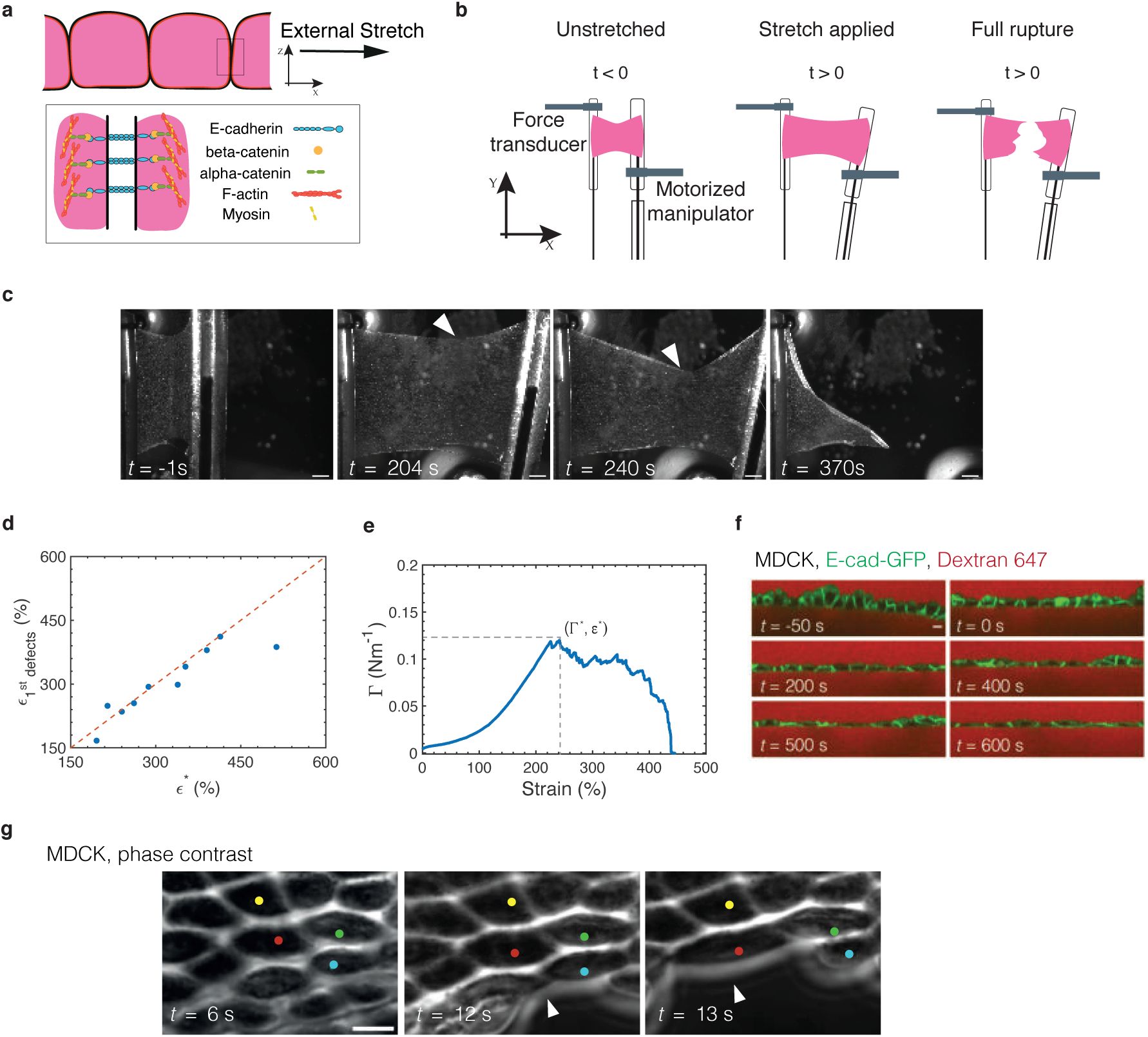
Epithelial monolayers rupture in response to excessive stretch. **(a)** Cellular-scale diagram of the epithelial monolayer. Top: profile view of the monolayer. Cells are linked to one another via specialised junctions. Bottom: zoom of an adherens junctions linking the F-actin cytoskeleton of neighbouring cells. The ectodomain of E-cadherin links cells to one another while its intracellular domain binds to the F-actin cytoskeleton via beta- and alpha-catenin. Myosin motor proteins bind F-actin to generate a cellular surface tension which results in a pre-tension in the monolayer. **(b)** Diagram of the experiment. Monolayers in pink are subjected to a ramp in deformation applied via displacement of one of the test rods. Stretch starts at time 0 and continues at a constant rate until full rupture of the monolayer. **(c)** Bright-field microscopy time series of an MDCK monolayer subjected to a ramp in deformation performed at 1% s^−1^. Arrowheads indicate the crack tip. Time is indicated in the bottom left corner. Scale bar = 500 µm. **(d)** Strain at which the first defects 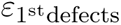 are observed as a function of the strain at which the maximum tension is reached *ɛ*^∗^. The dashed red line shows the line of slope 1. **(e)** Evolution of monolayer tension as a function of applied strain for the monolayer shown in c. Dashed lines show the maximum tension Γ^∗^ and strain *ɛ*^∗^ coinciding with the appearance of the first defects. Full rupture of the monolayer takes place for *ɛ*=450%. **(f)** Time series of a profile view of an MDCK monolayer during stretch. Intercellular junctions are visualised with E-cadherin-GFP (green) and cells are visualised by dye exclusion of Dextran-Alexa647 (red) added to the medium. Time is indicated in the bottom left corner of each image. Deformation starts at time 0 and proceeds at a rate of 1%/s. Scale bar = 10 µm. **(g)** High magnification phase contrast time series of crack propagation along cellular interfaces in an MDCK monolayer. The crack front is indicated by white arrowheads. Several cells are marked by coloured dots in each frame. Time is indicated in the bottom left corner. Scale bar = 10 µm.

After preconditioning (materials and methods), we subjected monolayers to a ramp in deformation applied at a constant strain rate of 1% s^−1^ (**Fig. 1b**, **Movie S1**) to minimise stress originating from viscoelastic contributions [20], and we monitored stress as a function of time. Monolayers could withstand more than a 3-fold increase in length before the first crack appeared and failed for strains of ∼ 300% (**Fig. 1c, e**). Most cracks first appeared at either of the free edges, although some could also be observed in the bulk of the material (**Fig. 1c**, **Fig. S1c**, **Fig. S2a**). Once nucleated, the crack front propagated through the material following a complex path alternating periods of rapid propagation and pauses until complete failure (**Fig. 1c** and **Fig. S1e**, **Movie S1**). As the monolayer length and width (in mm) are several orders of magnitude larger than its thickness (∼ 10 *µ*m), we approximated the tissue to a thin sheet and normalised the force *F* to the average width *w*_0_ of the monolayer prior to deformation (**Fig. S1d**) to generate tension-strain curves. In response to deformation, tension first rose linearly up to ∼ 50% strain before increasing more rapidly until reaching a peak tension Γ^∗^ = *F* ^∗^*/w*_0_ (**Fig. 1e** and **Fig. S1a-b**). Beyond this strain, cracks became apparent and tension decreased until complete failure of the monolayer.

In many materials, the peak tension marks the onset of fracture. Therefore, we plotted the strain *ε*^∗^ at which Γ^∗^ was reached as a function of the strain 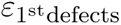 at which we visually observed the first monolayer defects in our time-lapse images (**Fig. 1c**). This revealed a clear correlation (**Fig. 1d**), suggesting that crack initiation results from accumulated stress in the tissue. Γ^∗^ did not depend on the dimensions of the monolayers, confirming that it represents a material property (**Fig. S1f**). Therefore, in the following, we used Γ^∗^ and *ε*^∗^ as parameters to characterise the onset of fracture in epithelial monolayers. As *ε*^∗^ was far greater than the strain for which monolayer unfurling is observed (up to ∼ 75%)[21], we considered monolayers to be two-dimensional sheets with a constant width in our analysis.

### Cracks occur at cell-cell junctions

After characterising fracture at the tissue-scale, we examined the phenomenon at the cell-scale. Cracks could in principle appear either because of cell lysis or detachment at intercellular junctions. High magnification phase contrast imaging revealed that cells were often very elongated at the free edge of the monolayer where most cracks initiate (**Fig. 1g**). As strain increased, cell-cell contacts appeared to progressively decrease in size perpendicular to the direction of applied stretch before the cells lost contact with one another. The crack front alternated phases of rapid propagation and pauses with no apparent preferred directionality, mirroring what was observed at the tissue-scale (**Fig. S1e**). In all cases, the crack front followed cell-cell junctions and cells appeared to peel apart from one another at intercellular contacts. When we examined the localisation of E-cadherin-GFP (a protein whose extracellular domains link cells to one another), we noticed disappearance of the cadherin signal from the cell surface after a cell-cell junction had ruptured (**Fig. S1c**, **Movie S2**). In profile views of monolayers, we observed a progressive decrease in the height of adherens junctions as strain increased (**Fig. 1f**). However, what happens to intercellular adhesion proteins at the molecular scale during change in intercellular junction height remains unclear.

### Epithelial monolayers can self-rupture by increasing their contractility

In vivo, epithelia can rupture in response to an increase in contractility. In some cases, this gives rise to defects that will impair the viability of the organism, while in others it forms part of the normal developmental program. For example, in the *Drosophila* amnioserosa, depletion of the myosin phosphatase MBS gives rise to small localised tears in the tissue disrupting the next developmental stage [17]. Conversely, during eversion of the *Drosophila* leg [12], the peripodial membrane must irreversibly rupture to allow correct development of the leg. To better understand the response of tissues to active stresses generated by myosins, we examined the response of epithelia to calyculin, a myosin phosphatase inhibitor that increases monolayer tension [22].

In our experiments, we incubated monolayers with 20 nM calyculin and monitored their response in tension while imaging their morphology (**Fig. 2a, f**, **Movie S3**). Monolayers were unperturbed until ∼ 90 minutes after the addition of calyculin, when holes appeared in the epithelium (**Fig. 2b, f**). These holes grew in time through complex crack propagation, merging together and eventually causing full rupture of the monolayer (**Fig. 2f**). As in ramp experiments, cracking occurred at intercellular junctions (**Fig. 2b**) and, although E-cadherin was lost from the cell surfaces whose intercellular junctions had ruptured (**Fig. 2b**), we did not notice any systematic loss of E-cadherin preceding crack formation. In contrast to ramp experiments, all cracks formed in the bulk of the tissue rather than at the edges (**Fig. 2f, S2d**), perhaps due to the isotropic nature of cortical contractility. From the onset of treatment, tension gradually rose in the monolayer reaching a peak after ∼ 90 minutes before decreasing as tissue fracture progressed (**Fig. 2c, S3a, c**). Similar to the ramp experiments, the time 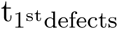 at which the first defect was observed was correlated with the time t^∗^ at which tissue tension reached a maximum, further confirming Γ^∗^ and t^∗^ as good criteria for characterising rupture onset (**Fig. 2d**). Intriguingly, tension at rupture was ∼ 10-fold lower than for ramps and the rupture time ∼ 10 times longer.

**Figure 2:**
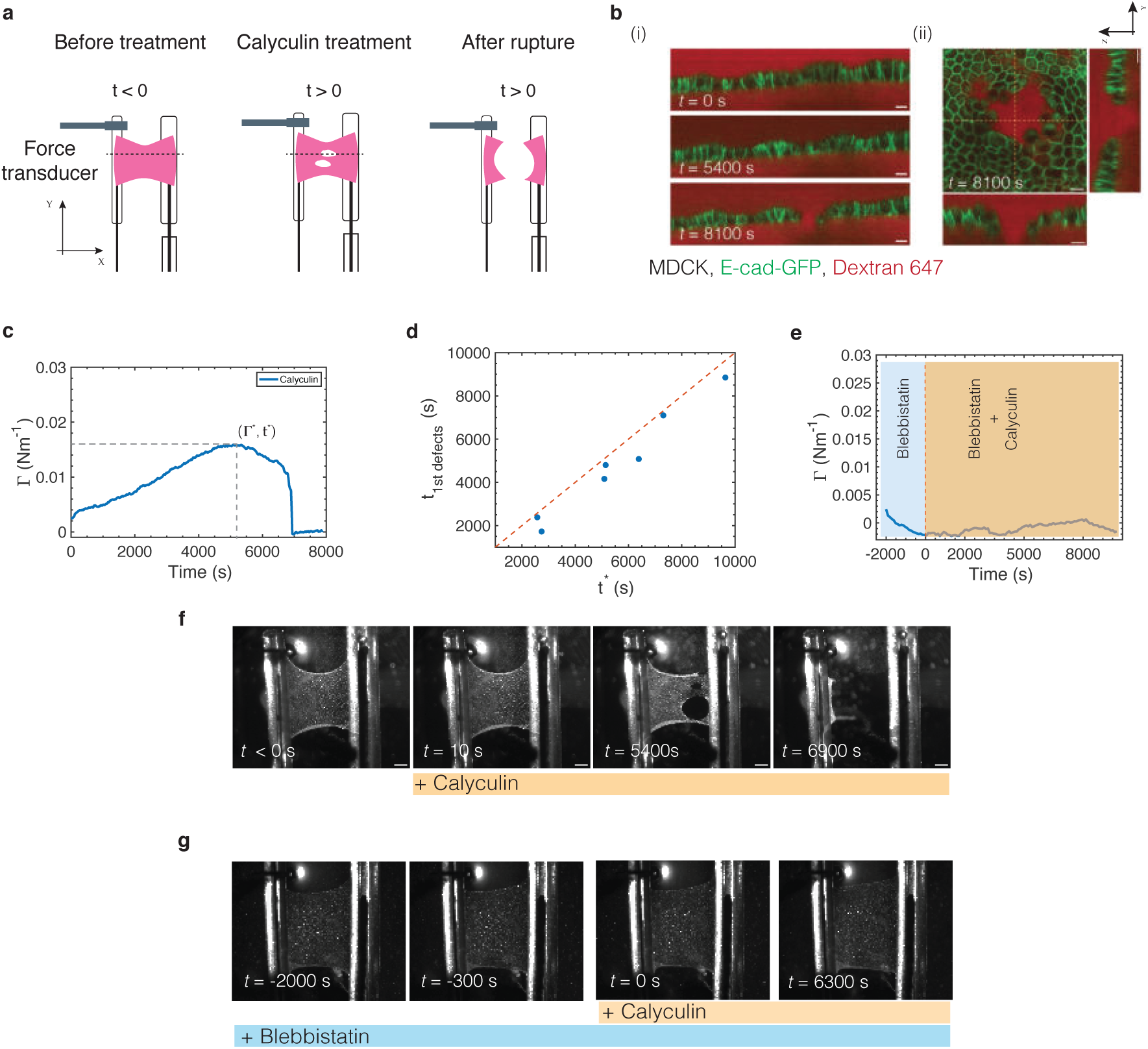
Monolayers can self rupture by increasing their myosin contractility. **(a)** Diagram of the experiment. Monolayers were treated with calyculin, an inhibitor of myosin phosphatase, at t=0. The tension in the monolayer was measured over time and the length of the monolayer was kept constant by the micromanipulator. After some time, defects appeared in the monolayer and measurements were continued until the monolayer failed. The dashed black line indicates a representative position at which the monolayer profile would be imaged. **(b)** Representative confocal images of a monolayer profile during calyculin treatment. Intercellular junctions are visualised with E-cadherin-GFP (green) and cells are visualised by dye exclusion of Dextran-Alexa647 (red) added to the medium. Scale bar = 10µm. (i) Profile view over time. Time is indicated in the bottom left corner of each image. (ii) XY view of a defect in a monolayer and its corresponding profile views in XZ and YZ. Dashed yellow lines indicate the position of the profile views. Time is indicated in the bottom left corner. (c) Temporal evolution of tension for the calyculin-treated monolayer shown in f. The dashed lines indicate the maximum tension Γ^∗^ and its timing *t*^∗^. **(d)** Time at which the first defects 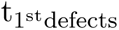 are observed as a function of the time at which the maximum tension is reached *t*^∗^. The dashed red line shows the line of slope 1. **(e)** Temporal evolution of tension for a monolayer treated with blebbistatin and calyculin, shown in g. The monolayer was first treated with blebbistatin alone for 2000s (blue shaded region) before calyculin was added (orange shaded region). **(f)** Bright field time series showing a representative calyculin-treated monolayer. Calyculin is added to the medium at time 0. Time is indicated in the bottom left corner. Temporal evolution of tension is shown in c. Scale bar= 500µm. **(g)** Bright field time series of a representative monolayer treated with blebbistatin and calyculin. Calyculin was added at time 0. Blebbistatin treatment was started at t= −2000s and was present throughout the experiment. Temporal evolution of tension is shown in e. Time is indicated in the bottom left corner. Scale bar= 500µm.

Given that calyculin is a broad spectrum phosphatase inhibitor, we confirmed that the monolayer rupture and the increase in tension that we observe are specific to cell contractility. First, using immunostaining, we verified that calyculin increased myosin phosphorylation but did not appear to perturb E-cadherin or cytokeratins over durations for which we typically observed changes in monolayer tension and rupture (**Fig. S3g - i**). Next, we incubated the monolayers with a specific myosin inhibitor, blebbistatin, for 30 - 40 min prior to calyculin addition (**Fig. 2g**, **Movie S4**). This ensured that any effect of calyculin specific to myosin contractility was inhibited. When only blebbistatin was present, tension decreased, as expected [22] (**Fig. 2e**, blue shaded area). After calyculin addition, the tension remained low and monolayers did not rupture over durations that typically led to failure when using calyculin alone (**Fig. 2e, g** and **Fig. S3b-f**, **Movie S5**). Altogether, we concluded that increasing myosin contractility in suspended epithelial monolayers is sufficient to generate rupture.

### Rupture strain, tension, and time scale with strain rate

During normal physiological function, cells in some adult tissues can experience deformations of hundreds of percent at strain rates ranging up to 100% s^−1^, while others experience far slower loading [1]. To explore this while minimising viscous contributions, we examined crack formation in monolayers subjected to ramps in deformation for a range of strain rates (0.1 - 3% s^−1^, Fig. S 4g).

For each monolayer, we characterised the rupture tension Γ^∗^ as well as the rupture strain *ε*^∗^ and the time t^∗^ at which Γ^∗^ was reached. When we plotted our data as a function of strain rate, we found that Γ^∗^ increased with strain rate, from 0.04 Nm^−1^ at 0.1% s^−1^ until seemingly reaching a plateau of ∼ 0.20 Nm^−1^ for strain rates above 2% s^−1^ (**Fig. 3a, S4a**). In contrast, both *ε*^∗^ and t^∗^ decreased with increasing strain rate. Rupture strain decreased from ∼ 700% at 0.1% s^−1^ until reaching a plateau of ∼ 250% for the larger strain rates, comparable to those observed in *T. adhaerens* which deforms at 20% s^−1^ [13] (**Fig. 3b, S4b**). Rupture time t^∗^ spanned nearly two orders of magnitude decreasing from 5.10^3^ to 10^2^ s (**Fig. 3c, S4c**). As previous work has shown that no cell rearrangements and only few cell divisions take place over hour-long periods in our suspended monolayers [23], these large deformations are likely accommodated through remodelling and deformation of the cytoskeleton and adhesive complexes. Interestingly, for the lowest strain rate 0.1% s^−1^, rupture tension Γ^∗^ and time t^∗^ were comparable to those observed in response to calyculin treatment, suggesting that rupture due to active stresses generated by myosin contractility and passive stresses due to deformation of the cytoskeleton may arise from the same biophysical processes.

**Figure 3:**
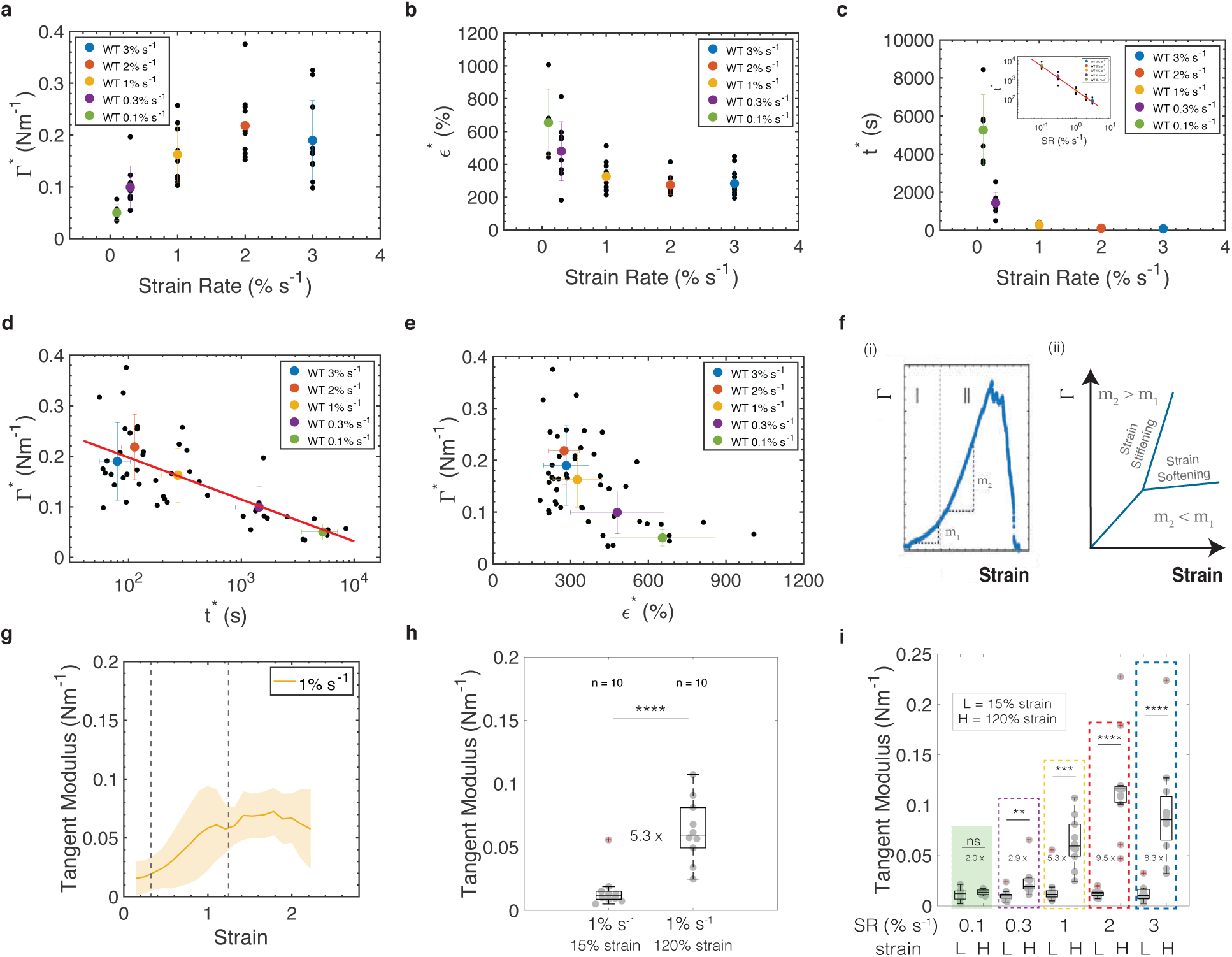
Rupture characteristics depend on strain rate. In all box plots, the central mark indicates the median, and the bottom and top edges of the box indicate the 25th and 75th percentiles, respectively. The whiskers extend to the most extreme data points that are not outliers. Data points appear as grey dots. Outliers are indicated with a red ‘+’ symbol. Statistically significant difference: ns non significant P *>* 0.05, *P *<* 0.05, ***P *<* 0.001, ****P *<* 0.0001, Kolmogorov-Smirnov test. Data was acquired from n = 6 monolayers for 0.1% s^−1^, n = 9 for 0.3% s^−1^, n = 10 for 1% s^−1^, n = 11 for 2% s^−1^, and n = 11 for 3% s^−1^. **(a)** Rupture tension, **(b)** rupture strain and **(c)** rupture time as functions of the strain rate. The inset in c shows the same data plotted on a log-log scale. **(d)** Rupture tension as a function of the rupture time. The graph is plotted on a semi-log scale because of the large range of time-scales associated with the different strain rates. The red line is a linear fit to the experimental data. **(e)** Rupture tension as a function of rupture strain. In panels (a-e) Black dots represent individual monolayers. Coloured dots represent the average value and whiskers represent the standard deviations. **(f)** Monolayers display a strain stiffening behaviour above 50% strain. **(i)** Diagram showing the change in slope in a typical tension-strain curve, characteristic of the strain stiffening behaviour observed in monolayers. The dashed line delimits the approximate change in regime. The slope m2 observed at higher strain (region II) is larger than that the slope m1 observed at lower strain (region I). (ii) Example showing the difference in the tension-strain curve that a material will display depending on whether it stiffens (slope increases) or it softens (slope decreases). **(g)** Tangent modulus as a function of strain for monolayers subjected to a ramp in deformation at 1% s^−1^. The average value is represented by the thick line and the shaded area shows the standard deviation. The dashed lines show the strains at which the tangent moduli were measured for comparison between low strain and high strain (n = 10). **(h)** Tangent modulus at low strain (*ɛ*= 15%) and at high strain (*ɛ*= 120%) for experiments pooled in panel (g). The fold change is indicated between the two strain magnitudes and 10 monolayers were examined. **(i)** Tangent modulus at low strain (L ≃ 15%) and at high strains (H ≃ 120%) as a function of strain rate. The fold change for each strain rate is indicated between the low and high strain regimes (SR 0.1%s^−1^ n = 6, SR 0.3%s^−1^ n = 9, SR 1%s^−1^ n = 10, SR 2%s^−1^ n = 11, SR 3%s^−1^ n = 11.

To gain insight into the failure mechanism of the material, we plotted the rupture tension as a function of the rupture time and rupture strain. This revealed that rupture tension Γ^∗^ scaled as ∼ log(1/t^∗^) (**Fig. 3d**), reminiscent of the failure dynamics of groups of microscopic bonds subjected to force [24, 25]. Experiments increasing contractility clustered close to the experimental data from 0.1% s^−1^ ramps (**Fig. 4a**). However, the rate of increase in tension dΓ^∗^/dt was several-fold larger in experiments involving deformation (**Fig. S4h-i**), suggesting some fundamental differences between these experimental conditions. In deformation experiments, rupture tension decreased linearly with rupture strain (**Fig. 3e**) and, as a consequence, the strain energy at rupture onset remained approximately constant across strain rates (**Fig. S4g**). The decrease of Γ^∗^ with *ε*^∗^ is surprising because our previous work suggested that monolayers behave as elastic solids for strain rates lower than 1% s^−1^ [20], and, in elastic solids, tension increases with strain, contrary to what is observed for rupture tension in our experiments (**Fig. 3e**). Such discrepancy may be due to the very large deformations used in the current study.

**Figure 4:**
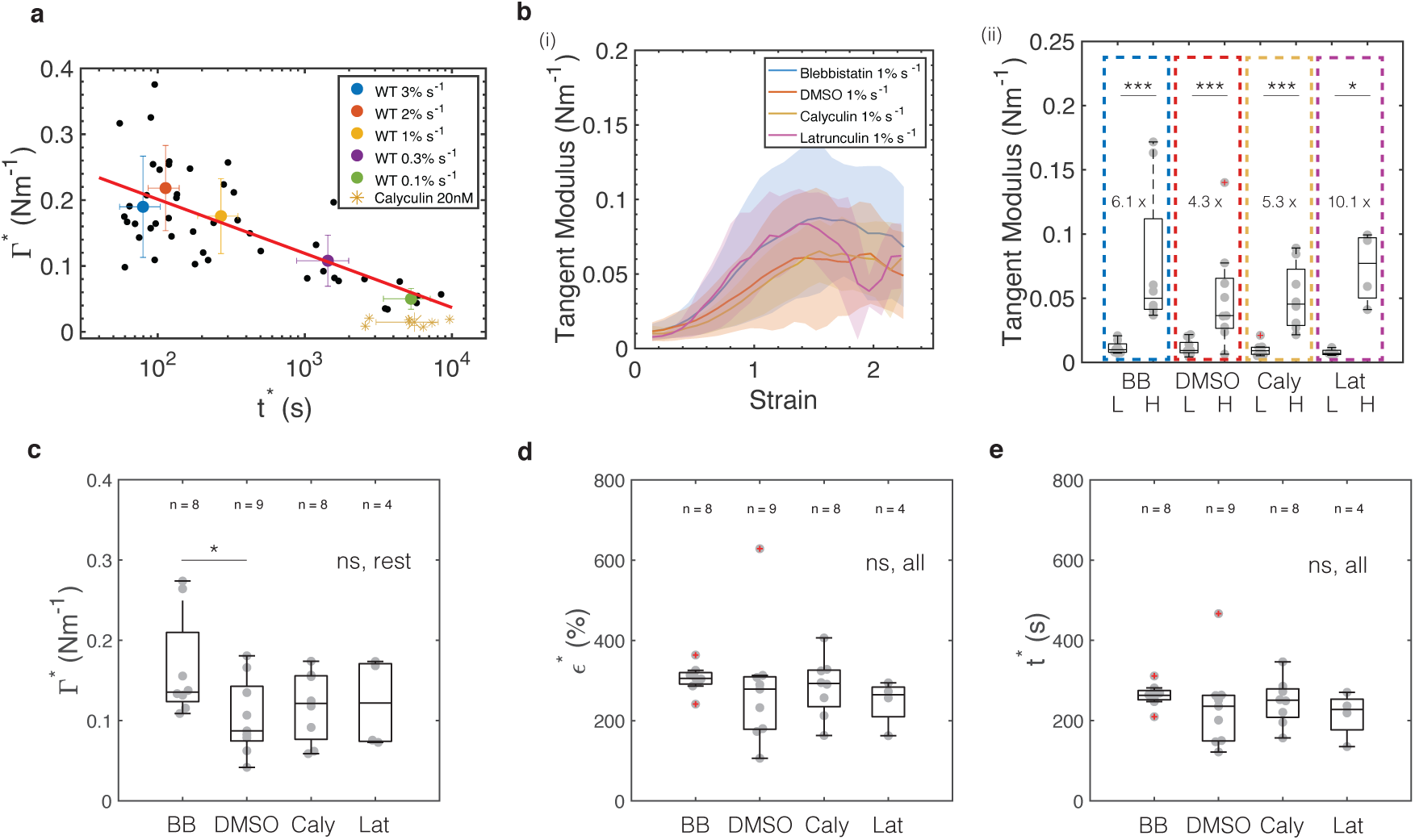
Strain stiffening does not depend on the actomyosin cytoskeleton. In all box plots, the central mark indicates the median, and the bottom and top edges of the box indicate the 25th and 75th percentiles, respectively. The whiskers extend to the most extreme data points that are not outliers. Data points appear as grey dots. Outliers are indicated with a red ‘+’ symbol. Statistically significant difference: ns non significant: P *>* 0.05, *:P *<* 0.05, ***:P *<* 0.001, ****:P *<* 0.0001, Kolmogorov-Smirnov test. Data was acquired from n = 8 monolayers for blebbistatin, n = 9 for DMSO, n = 8 for calyculin, and n = 4 for latrunculin. **(a)** Semi-log graph of the rupture tension as a function of the rupture time for all strain rates and for calyculin-treated monolayers. Black dots represent individual monolayers, except for calyculin treated monolayers that are represented by yellow asterisks. Coloured dots represent the average value and whiskers represent the standard deviations, except for calyculin treated monolayers that are represented by a cross. Note that calyculin treated monolayers cluster very close to the trend line expected for experiments in which strain rate is varied. **(b)** Tangent modulus for monolayers in which actomyosin has been perturbed. **(i)** Average tangent modulus (thick lines) and standard deviation (shaded area) as a function of strain for ramp experiments performed at 0.3% s^−1^ for monolayers pre-incubated with blebbistatin 50µM (BB, blue, n = 8), DMSO (red, n = 9), calyculin 20 nM (Caly, yellow, n = 8) and latrunculin 1µM (Lat, purple, n = 4). (ii) Boxplots comparing the tension gradient at low strain (L ≃ 15% strain) and high strain (H ≃ 120 % strain) for monolayers treated with drugs perturbing the actomyosin cytoskeleton. (c-e) Boxplots showing the **(c)** rupture tension, **(d)** rupture strain, and **(e)** rupture time for monolayers pre-incubated with different treatments perturbing the actomyosin cytoskeleton.

### Monolayers display a strain rate-dependent strain stiffening

One potential origin for this counter-intuitive behaviour could involve a change in the mechanical response of the monolayer with strain rate. In monolayers stretched at 1% s^−1^, the slope of the tension-strain curve visibly increased for strains larger than ∼ 50% (**Fig. 1e**). This feature, known as strain-stiffening (**Fig. 3f**), has also been observed in biopolymer networks, allowing them to limit deformation [26]. Therefore, we characterised strain stiffening further and examined how it changed with strain rate. When we computed the gradient of the tension-strain curve (i.e. the tangent modulus) for monolayers subjected to ramps at strain rates of 1% s^−1^, we observed three distinct regimes: first, until ∼ 30% strain, the tangent modulus was constant; then, between ∼ 50% and 100% it increased monotonically with strain, and finally from around 100% strain, it reached a plateau (**Fig. 3g**). The tangent modulus in the third regime was ∼ 5-fold larger than in the first (**Fig. 3h**). Intriguingly, monolayer strain-stiffening was dependent on strain rate (**Fig. 3i**). At the lowest strain rate (0.1% s^−1^), no change in the tangent modulus at high strain was apparent but, from 0.3% s ^−1^, it increased with strain rate, saturating for rates above 2% s^−1^, a behaviour known as shear-stiffening (**Fig. S4e**, f) [27, 28, 29, 30].

Strain stiffening might arise from mechanotransductory processes or from the intrinsic organisation of the cytoskeleton and cells within monolayers. Stiffening occurred for strain rates above 0.3% s^−1^ with the most pronounced observed for 3% s^−1^, signifying that a hypothetical mechanotransductory process should take place over durations shorter than ∼ 300 seconds. This points to adaptive processes involving post-translational modifications or protein recruitment downstream of detection rather than transcriptional changes. Therefore, we examined known mechanisms such as recruitment of vinculin downstream of the unfolding of alpha-catenin [31, 32], as well as tension-induced recruitment of EPLIN [33] and E-cadherin [34]. For this, we generated monolayers of cells expressing each of these proteins tagged with GFP and subjected them to strains of 30 - 80% for 30 minutes. However, we could not observe recruitment of any of these candidates (**Fig. S5**). This suggested that strain stiffening is more likely to arise from an intrinsic response of the cytoskeleton in monolayers. We therefore decided to investigate the role of the main cytoskeletal components (actomyosin and intermediate filaments) in strain stiffening and how strain stiffening affects the strength of epithelial tissues.

### Strain-stiffening and fracture are independent of actomyosin

Actomyosin is arguably the main determinant of cell shape and a key player in cell mechanics. In tissues subjected to low strain (less than 30%, before strain stiffening is observed), the actomyosin cortex controls monolayer rheology [20] and increases in cortical contractility lead to stress stiffening [22]. Therefore, we examined the role of myosin in strain stiffening by repeating our ramp deformation experiments at 1% s^−1^ for monolayers in which myosin contractility had been inhibited or enhanced (**Fig. 4b, S6d**). For this, we blocked myosin activity with blebbistatin, and we increased phospho-myosin levels by treating monolayers with calyculin for 20 minutes, a duration sufficient to increase pre-tension Γ_0_ significantly but not to cause rupture (**Fig. 2c, Movies S6, S8**). Neither treatment affected strain stiffening (**Fig. 4b**). Next, to examine the role of F-actin, we treated monolayers with latrunculin, an inhibitor of actin polymerisation that leads to global loss of F-actin (**Fig. S6f**, **Movie S9**). Similar to myosin, F-actin appeared to play no role in strain-stiffening (**Fig. 4b, S6d**). Interestingly, none of these treatments affected the rupture characteristics of monolayers (**Fig. 4c - e, S6a - c**). Hence, the strength of epithelia appears independent of actomyosin.

### Keratin filaments control monolayer strength

We then focused our attention on keratin intermediate filaments. These form entangled networks in the vicinity of the nucleus with wavy filament bundles radiating out towards the cell periphery where they connect to neighbouring cells via specialised protein complexes known as desmosomes. These comprise desmosomal cadherins whose extracellular domains bind to counterparts on adjacent cells while their cytoplasmic domains connect to keratin filaments via ancillary proteins such as desmoplakin, plakoglobin, and plakophilin [35]. Keratins can extend to multiple times their original length before rupture as well as bear high tensile load and keratin networks strain stiffen in vitro [36, 37], making keratins good candidates to maintain tissue integrity and control monolayer rheology at high strain [38, 39, 40, 41]. Previous work has highlighted a mechanical role for desmosomes and shown that, as epithelia are stretched, desmosomes become progressively loaded [42]. Keratins and desmosomes have also received much interest from a clinical standpoint for their role in the maintenance of skin integrity. Indeed, mutations in keratins and desmosomal proteins give rise to skin blistering disorders with symptoms of mechanical fragility [43, 5]. Thus, the supracellular network formed by keratins and desmosomes appears as a promising structure to contribute to tissue strength.

We first imaged the keratin filament network conformation in response to stretch by imaging monolayers stably expressing K18-GFP at high magnification. At low strain, keratin bundles extending towards the cell periphery displayed wavy morphologies, similar to those observed on glass (**Fig. 5a**). As strain progressively increased, the network became visibly stretched in the direction of deformation, suggesting that keratin filament bundles become loaded as previously reported [38, 44]. Even when applying strains larger than 300%, the network retained its structural integrity, although defects did appear above 400% strain.

**Figure 5:**
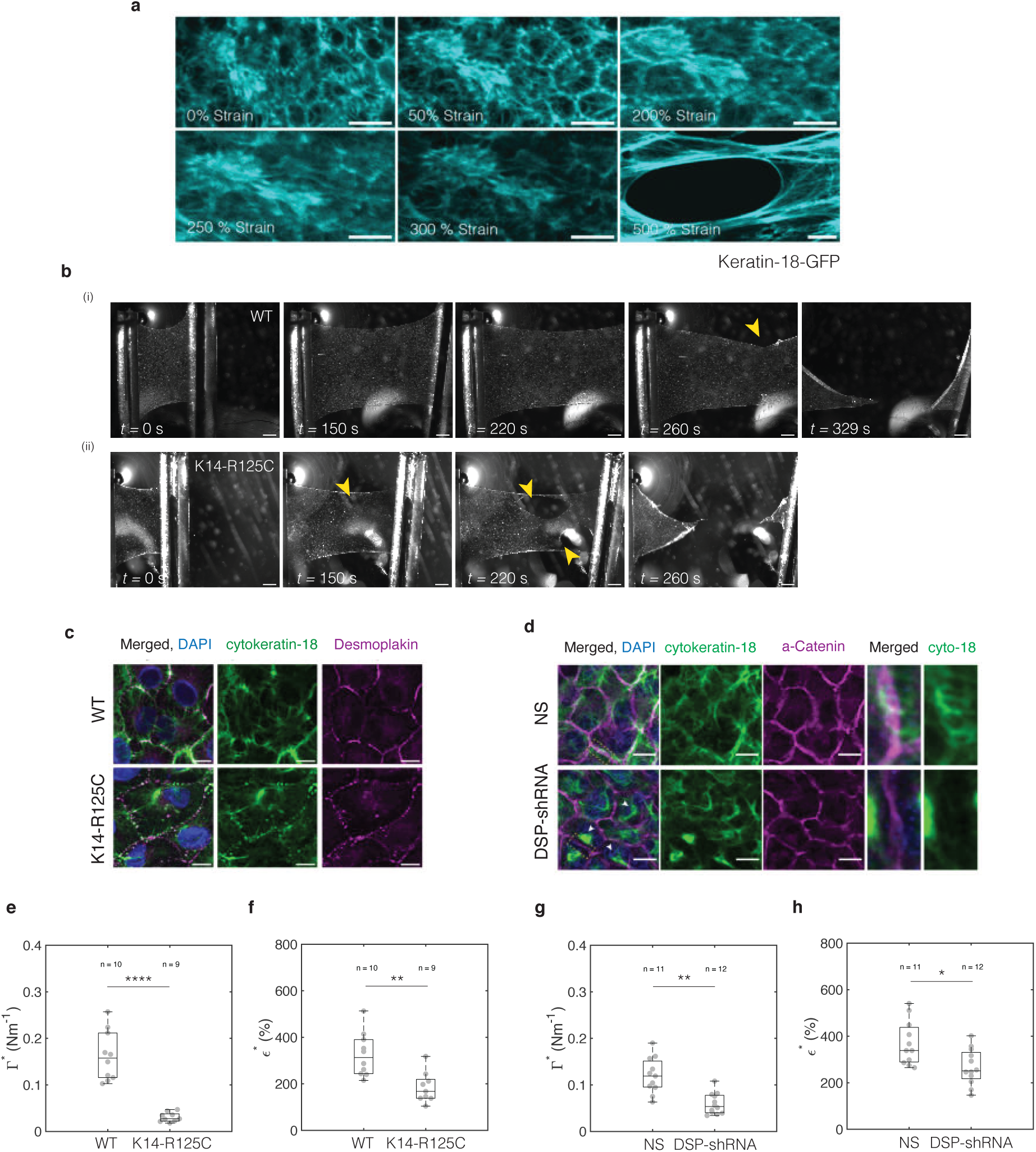
Perturbation of the keratin intermediate filament network fragilises monolayers. **(a)** Image series of Keratin-18-GFP localisation in cells in suspended monolayers subjected to increasing deformation. The strain is indicated in the bottom left corner. Scale bar= 10µm. **(b)** Bright-field time series of (i) representative wild-type (WT) and (ii) K14, R125C monolayers during a ramp experiment performed at 1% s^−1^. Arrowheads indicate the start and growth of cracks. Time is indicated in the bottom left corner. Scale bar= 500 µm. **(c)** Immunostaining of cytokeratin-18 (green) and desmoplakin (magenta) in WT (top row) and K14, R125C monolayers (bottom row). Scale bar= 10µm. **(d)** Immunostaining of cytokeratin-18 (green) and desmoplakinalpha-catenin (magenta) in non-silencing shRNA (NS, top row) and desmoplakin-shRNA monolayers (DSP-shRNA, bottom row). Scale bar= 10µm. The 4th and 5th column show a zoom of an intercellular junction. **(e-h)** In all box plots, the central mark indicates the median, and the bottom and top edges of the box indicate the 25th and 75th percentiles, respectively. The whiskers extend to the most extreme data points that are not outliers. Data points appear as grey dots. Statistically significant difference: ns non significant: P *>* 0.05, *:P *<* 0.05, ***:P *<* 0.001, ****:P *<* 0.0001, Kolmogorov-Smirnov-test. **(e-f)** Box plots comparing the **(e)** rupture tension and **(f)** rupture strain and between WT and K14, R125C monolayers. **(g-h)** Box plots comparing the **(g)** rupture tension and **(h)** rupture strain between non-silencing shRNA (NS) and desmoplakin-shRNA (DSP-shRNA) monolayers.

To understand the role of keratins and desmosomes, we perturbed each of these in turn and assessed their impact on tissue mechanics. To study the contribution of keratins, we overexpressed a dominant mutant of keratin 14, K14-R125C, a mutation identified in some epidermolysis bullosa patients that fragilises keratin filaments and causes disaggregation of the network [43, 45, 46]. Immunostaining against keratin 18 in K14-R125C monolayers revealed a highly disrupted keratin network compared to the extensive supracellular network observed in control monolayers, consistent with previous reports [47, 48] (**Fig. 5c**). Although the desmoplakin organisation appeared normal, connections of intermediate filaments to desmosomes were missing. To determine if the connection between cellular keratin networks is important for tissue mechanics and force transmission, we also engineered a cell line with a stable desmoplakin depletion (DSP-shRNA) to perturb desmosomes (**Fig. S7**). In DSP-shRNA cells, the keratin network failed to attach to intercellular junctions (**Fig. 5d**, zoomed region) and it often appeared collapsed around the nucleus rather than displaying the characteristic stellate organisation observed in wild-type cells (**Fig. 5d**, arrowheads).

We then examined whether these perturbations affected tissue mechanical response by performing ramp experiments at 1% s^−1^ (**Fig. 5b, S8a, Movies S1, S10** and **Movies S11, S12**). Quaitatively, both perturbations led to earlier rupture than in controls (**Fig. 5b, S8a** arrowheads). Neither crack propagation velocity (**Fig. S8j**) nor the location of crack nucleation were affected by perturbation of the keratin network (**Fig. S2c**). Quantitatively, both the rupture strain, *ε*^∗^, and the rupture tension, Γ^∗^, were significantly lower in the perturbed tissues (**Fig. 5e-h**, **S8f-i**). These data confirm that keratin networks connected across cells are necessary for monolayers to resist large deformations and that their absence leads to more fragile tissues.

### Strain stiffening depends on the integrity of a keratin supracellular network

We next investigated the contribution of keratin filament networks to strain-stiffening by computing the tangent modulus in K14-R125C and DSP-shRNA monolayers subjected to ramps at 1% s^−1^ (**Fig. 6a-d**). These presented two main features that differed from controls. First, in both perturbations, there was no increase in the tangent modulus with strain in contrast to controls (**Fig. 6b, d**). This indicated that both the keratin network and its attachment to cell junctions are necessary for monolayers to strain-stiffen. Second, in the DSP-shRNA monolayers, we detected larger values of the tangent modulus at low strains compared to control (**Fig. 6d, S8d**). Although this was surprising, we reasoned this may arise because of a compensatory mechanism. For example, when tight junction proteins ZO-1 and ZO-2 are depleted, cells become more contractile[49].

**Figure 6:**
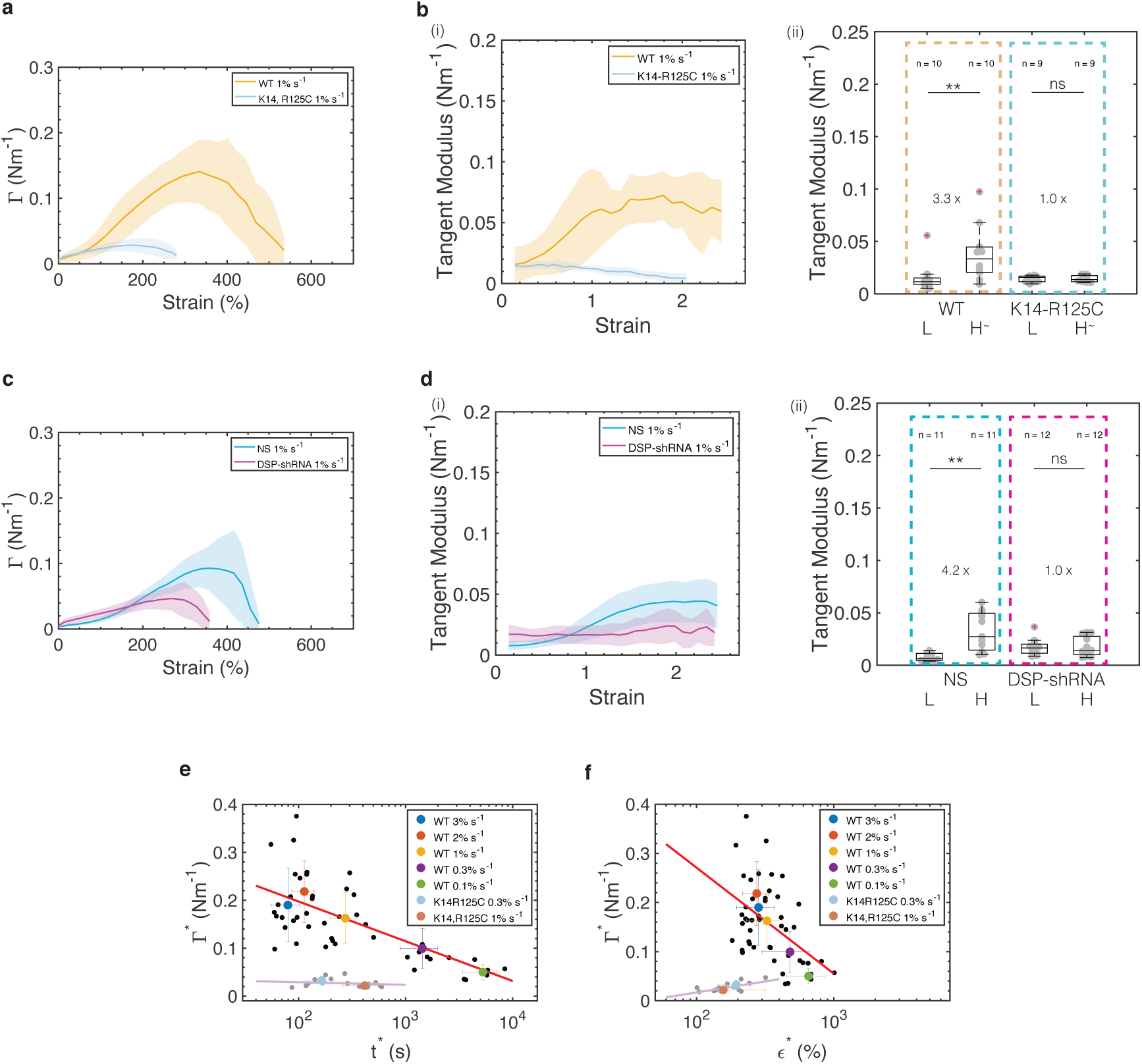
Keratin intermediate filament networks control tissue strain stiffening and strength. In all box plots, the central mark indicates the median, and the bottom and top edges of the box indicate the 25th and 75th percentiles, respectively. The whiskers extend to the most extreme data points that are not outliers. Data points appear as grey dots. Outliers are indicated with a red ‘+’ symbol. Statistically significant difference, ns non significant: P *>* 0.05, *:P *<* 0.05, ***:P *<* 0.001, ****:P *<* 0.0001, Kolmogorov-Smirnov-test. Data from n=10 WT, n=9 K14-R125C, n=11 non-silencing shRNA, and n=12 desmoplakin shRNA monolayers. **(a)** Tension as a function of strain for monolayers subjected to a ramp in deformation at 1% s^−1^ for WT (orange) and K14, R125C (blue) monolayers. Solid lines represent the average value and shaded areas show the standard deviation of the distribution. **(b)** (i) Tangent modulus as a function of strain for monolayers subjected to a ramp in deformation at 1% s^−1^ for WT (orange) and K14, R125C (blue) monolayers. Solid lines represent the average value and shaded areas show the standard deviation of the distribution. (ii) Box plots of the tangent modulus at low (L ≃ 15% strain) and high strain (H^∼^ ≃ 70% strain) in WT and K14, R125C monolayers. The fold change between low and high strains is indicated in between the box plots. **(c)** Tension as a function of strain for monolayers subjected to a ramp in deformation at 1% s^−1^ for non-silencing shRNA (NS, blue) and desmoplakin-shRNA (DSP-shRNA, magenta) monolayers. Solid lines represent the average value and shaded areas show the standard deviation of the distribution. **(d)** (i) Tangent modulus as a function of strain for monolayers subjected to a ramp in deformation at 1% s^−1^ for non-silencing shRNA (NS, blue) and desmoplakin-shRNA (DSP-shRNA, magenta). Solid lines represent the average value and shaded areas show the standard deviation of the distribution. (ii) Box plots of the tangent modulus at low (L ≃ 15% strain) and high strain (H≃ 120% strain) in NS-shRNA and DSP-shRNA monolayers. The fold change between low and high strains is indicated in between the box plots. (e-f) Data points are acquired for different strain rates. Black dots show indiviudal WT monolayers and grey dots individual K14, R125C monolayers. Coloured dots show the population average for a given strain rate. Whiskers indicate the standard deviation. The red and pink lines are linear regressions to the data. **(e)** Rupture tension as a function of rupture time. **(f)** Rupture tension as a function of rupture strain.

To investigate the molecular mechanism underlying the increase in tangent modulus, we immunostained DSP-shRNA and control monolayers against phospho-myosin (pMLC), F-actin, and E-cadherin. In DSP-shRNA monolayers, we visually observed an increase in pMLC and F-actin as well as E-cadherin (**Fig. S8e, S9g** respectively). This suggests that an increase in myosin contractility underlies the higher tangent modulus of DSP-shRNA monolayers at low strain, consistent with the stress-stiffening behaviour observed at low strain in response to short duration calyculin treatment reported in our previous work [22]. However, DSP-shRNA monolayers were more fragile at high strain despite having more F-actin, pMLC and E-cadherin (**Fig. S8e**, **Fig. S9g**).

When we perturbed actomyosin organisation, we could not observe any impact on strain stiffening (**Fig. 4b**) or the rupture characteristics of monolayers (**Fig. 4c - e**), implying that the structure that controls these properties is not affected by treatment with latrunculin, blebbistatin or calyculin. Therefore, we verified by immunostaining that these drugs did not affect the keratin network (**Fig. S3i, S6e**). Overall these results confirm that strain-stiffening is dependent on the presence of a supracellular keratin network, pointing to its key role in governing the mechanical response of monolayers at high strains.

Finally, we investigated if intermediate filament perturbation affected the strain rate dependency of rupture tension and strain (**Fig. S9a - d**). For this, we compared K14-R125C monolayers subjected to ramps at two different strain rates, 1% s^−1^ and 0.3% s^−1^. Neither rupture tension nor rupture strain changed with strain rate (**Fig. S9a**, b), in contrast to control monolayers (**Fig. 3a, b**). When we plotted these experiments in rupture tension-rupture strain or rupture tension-rupture time plots, we found that they presented a trend very different from the ramp experiments performed on WT monolayers (**Fig. 6e, f**).

In summary, we show that keratin intermediate filaments are primarily responsible for load bearing at high strain and that strain-stiffening also depends on this network. Altogether, our results point to keratin intermediate filaments as the main structure protecting epithelial monolayers against large deformations.

### A supracellular keratin filament network prevents rupture in developing epithelia

To determine the importance of keratin networks and desmosomes in physiological conditions, we examined their role in the epidermis of *Xenopus laevis* during body axis elongation, a process occurring at a very low strain rate driven by long range tensile forces [50, 51, 52]. We first characterised strain in wild type embryos to determine regions exposed to high strain, which may be affected by perturbation to keratins and desmosomes. For this, we microinjected fluorescent markers to visualise the membrane and the nucleus of epidermal cells by live microscopy. Our temporal analyses of cell morphology revealed that lateral epidermal cells undergo large deformation along the anteroposterior axis (**Fig. S10a**, b, d; **Movie S13**). After identifying the relevant desmosomal and keratin proteins expressed in the elongating lateral epidermis (**Fig. S11**), we analysed the impact of depleting desmoplakin and keratin 8 by microinjecting anti-sense oligonucleotides previously validated in *Xenopus laevis* [53, 54]. Cells in control embryos became visibly elongated in the anteroposterior axis between the beginning and end of body axis elongation (**Fig. 10b**), leading to cells with a significantly larger aspect ratio at late stages (**Fig. S10d**). Over the course of body axis elongation, average cell strain increased at a constant rate, reaching ∼40% by the end of this process (**Fig. S10e**). Whereas a clear keratin network was visible within the cells of wild-type embryos (**Fig. S10f**), it became less well defined and no longer appeared connected across cells in knockdown embryos (**Fig. S10f**, compare the membrane and the cytokeratin localisations). Morphologically, cells within knockdown tissues had a significantly lower aspect ratio than wild-type cells at the same stage (**Fig. S10d**), perhaps because the keratin network no longer transmitted long range tensile forces. Consistent with this idea, cell strain remained low over the course of body axis elongation (**Fig. S10e**, **Movie S14** middle). When the keratin supracellular network was perturbed, cells often detached from their neighbours creating small defects and these occasionally gave rise to larger cracks (**Fig. S10c**, **Movie S14** right). Crack area remained much smaller than in suspended monolayers, likely because of adhesion to underlying cell layers limits crack growth. Although knockdown embryos developed until late neurula stages, they died at later larval stages. In summary, a supracellular network of keratins linked by desmosomes is critical during development but the presence of additional boundary conditions on the basal side may limit crack growth.

### Non-linear rheology and collective dynamics of bond rupture control the strength of epithelial monolayers

The onset of rupture is associated with the separation of cell junctions. At the molecular level, this implies unstable dynamics with bonds dissociating more frequently than they associate. Simple models of interfaces linked by dynamic bonds have already demonstrated that a molecular slip bond behaviour may lead to a finite time of separation that decreases with the mechanical tension applied to the system [24, 25]. Such a trade-off between time-to-fracture and applied mechanical force is consistent with our observation that monolayers reach higher rupture tension Γ^∗^ and shorter rupture times t^∗^ at large strain rates (**Fig. 3a, c**). However, we also observed that the strain at the onset of fracture decreased with strain rate (**Fig. 3b**): the higher the rupture tension, the lower the strain at rupture (**Fig. 3e**). Remarkably, this qualitative trend is not present when the intermediate filament network is disrupted and the shear-stiffening behaviour of the monolayer is abrogated (**Fig. 6e**). Although separation of a junction is primarily controlled by the tension it is subjected to, this tension results from a multitude of processes, ranging from externally imposed deformations or forces due to active contractile behaviours. Furthermore, the boundary conditions influence the resulting mechanical state of the tissue. The onset of monolayer rupture therefore likely involves the interplay between the tissue-scale monolayer rheology and the molecular-scale dynamics of bonds. We hypothesise here that this interplay is responsible for the range of behaviours reported in this study.

A stochastic slip bond dynamics model inspired by the work of [24] and [25] combined with a range of material models enabled us to explore the qualitative and quantitative agreement of this hypothesis with our data (see methods). In essence, two surfaces representing an intercellular junction are connected by a population of *N* independent linkers that can exist in two states, bound or unbound (**Fig. 7a**). They bind at a fixed rate *k_on_*, but unbind at a rate *k_off_* that increases exponentially with mechanical load with a scale *f*_0_. In our implementation, all bound links bear an equal share of the tension applied to the surfaces. Rupture is assumed to be triggered when the system transitions to a state where all links detach (**Fig. 7b**). When a constant force is applied to a junction, we find as expected that the larger the applied force is, the shorter the time to rupture (**Fig. 7c**).

**Figure 7:**
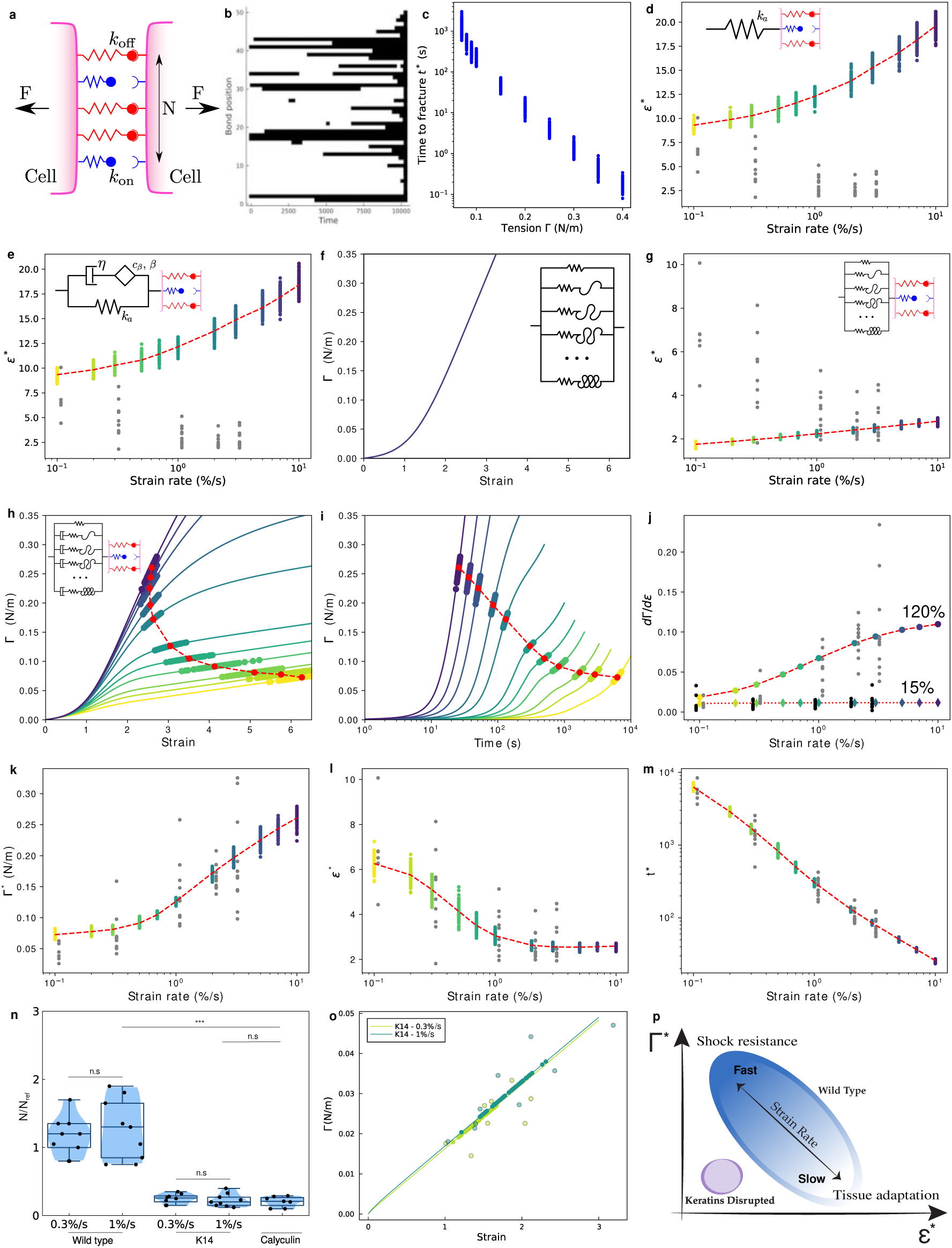
Multiscale modelling of rupture onset. **(d-e, g-m)** Each coloured dot represents a simulation run. 100 simulations were run for each strain rate. Each grey dot represents an experimental data point. Red dots indicate the mean value for a series of simulations. The dashed red lines link the mean value for each strain, strain rate, or time to show the trend. **(a)** Cell surfaces are linked by a population of N independent linkers with slip bond dynamics and subjected to a force F. Each linker can associate with its counterpart on the opposite surface with a rate constant *k_on_* and dissociate with a rate constant *k_off_*. The latter depends on the force *f* it is subjected to: *k_off_* = *k_off,_*_0_*e^f/f^*^0^ where *f*_0_ is a model parameter. The total force applied to the surface is assumed equally shared between closed bonds. **(b)** Typical evolution of the states of a population of 50 links over time when the junction is subjected to a constant tension. The black color indicates an unbound state. The simulation ends when all links are unbound, defining *t* = *t*^∗^. **(c)** Time to fracture *t*^∗^ as a function of the mechanical tension for junctions subjected to a constant tension. **(d)** Strain at rupture as a function of the ramp strain rate for a linear elastic material. The inset represents the rheological model used. **(e)** Strain at rupture as a function of the ramp strain rate for a linear visco-elastic material. The rheological model is shown in inset and is fitted to the rheology of wild-type MDCK monolayers at low strain [55]. **(f)** Tension-strain relationship for a strain stiffening material. The rheological model is shown in inset and it accounts for the strain stiffening behaviour observed at large strains in monolayers. **(g)** Strain at rupture as a function of the ramp strain rate for a strain stiffening material. The rheological model is shown in inset. **(h)** Tension-strain, and **(i)** tension-time characteristics (solid lines) for a shear stiffening material. The rheological model is presented as inset in h. Each curve represents a different strain rate. The colour code indicates the strain-rate consistently with panels j-m. Dark blue colours indicate large strain rates, while yellow colours indicate small strain rates as shown on panels d, e, and g. **(j)** Tangent modulus at 15% and 120% strain as a function of strain rate for a shear stiffening material. The rheological model is presented as an inset in h. Grey dots show experimental data at 120% and black dots at 15% strain. (k - m) Graphs showing the **(k)** tension, **(l)** strain, and **(m)** time at rupture as a function of the ramp strain rate for the model presented in h. **(n)** Normalised bond numbers required to fit the K14-R125C and calyculin treated samples in comparison to the WT monolayers. **(o)** Tension as a function of strain for the linear viscoelastic model presented in panel e, adjusted in stiffness to match the experimental stress values (*k_a_* = 1600 Pa instead of 760 Pa). The contact model parameters for K14-R125C mutants from n are used to simulate the rupture tension distributions at loading rates of 0.3%/s and 1%/s (filled green circles on the curves). Opn circles outside of the curves represent the experimental rupture tension and strain data for K14-R125C monolayers. **(p)** Summary diagram showing the range of behaviours displayed by monolayers. Wild-type monolayers (blue region) subjected to ramps at high strain rates have a high rupture tension and low rupture strain, making them an ideal material to resist shocks. At low strain rates, the rupture strain is large and the rupture stress is low, allowing the material to adapt. Disruption of keratin intermediate filaments abrogates these strain rate-dependent behaviours.

In our experiments, the force is not constant but instead results from ramps of deformation applied at different rates. The tension in the tissue for a given strain rate is a function of time that depends on the material’s rheology. For a range of strain rates and material models, we can calculate the temporal evolution of the applied tension and the bond dynamics can then be simulated a large number of times to get statistics for the rupture parameters Γ^∗^ and t^∗^. The corresponding value of *ε*^∗^ is then determined using the rheological model.

To grasp the role of rheology in the fracture behaviour, we first considered the case of a linear elastic relationship between stress and strain, which corresponds to the low strain rate limit identified in [20, 55] for relatively small deformations (less than 30%). As expected, the rupture load increases with strain rate (**Fig. 12a**), but so does the strain at rupture (**Fig. 7d**); this arises because stress and strain are proportionally related to each other through the constitutive equation of the material. A more complex material model takes into account the rich time-dependent linear rheological behaviour that we previously validated at low deformation [55] and that is controlled by actomyosin [20] (**Fig. 12b**). However, even in this case, the relationship between rupture strain and strain rate remains a monotonically increasing function (**Fig. 7e**).

Our experimental observations show that the material strain-stiffens for strains larger than ∼ 50%, but also that stiffening is larger for large strain rates. Strain stiffening is a classical feature of random fibre networks [56, 57, 58]. The phenomenon can be understood in a number of ways; at a basic level, as strain increases, more and more fibres align along the direction of deformation and become taut, signifying that more fibres are progressively recruited to carry the mechanical load. As the stiffness of the network increases with the number of recruited fibres, it naturally increases with applied strain. In our case, intermediate filament bundles appear responsible for both strain-stiffening and shear-stiffening (**Fig. 6**). As keratin intermediate filaments do not intrinsically strain-stiffen [41], we hypothesise that progressive recruitment of intermediate filaments to bear load underlies the strain stiffening observed in experiments. To mimic this, we introduce a non-linear elastic model where springs are progressively recruited as the deformation increases (inset, **Fig. 7f**), leading to the tension-strain characteristics presented in **Fig. 7f** (see methods). This implementation displays gradients consistent with the experimental trend at 15% and 120% strain at high strain rate, recapitulating the strain stiffening behaviour in this regime. The model can be calibrated to provide a good agreement with the experimental stress at high strain rates (**Fig. 12c**). However, when it is combined with the bond model, strain stiffening alone fails nonetheless to capture the qualitative relationship between rupture strain and strain rate (**Fig. 7g**). This is to be expected as the lack of time-dependent behaviour implies that the rupture strain necessarily increases with the rupture stress, which increases with strain rate.

The strain-rate dependent strain-stiffening behaviour of the network (or shear stiffening), therefore, appears necessary to account for the inverse correlation between tension and strain at rupture. A possible origin for such a behaviour is that the keratin bundle network relaxes tension in a visco-elastic manner. Whenever a bundle is recruited as a result of the stretch, it would then behave not as an elastic component but as a viscoelastic Maxwell-like fluid, as a first order approximation. If we stretch faster than the relaxation time-scale of this system, all filaments can be recruited and the strain-stiffening is visible; whereas if we stretch slower, relaxation occurs faster than the rate at which new filaments are recruited and the stiffening behaviour is not observed. To implement this, we considered a material model consisting of many Maxwell branches in parallel, all identical but becoming load-bearing at different strain thresholds (**Fig. 7h**, inset), consistently with the strain-stiffening model caligrated at high rate. This non-linear visco-elastic model reproduces the shear-stiffening behaviour observed in experiments (**Fig. 7h - j**). After calibration, the combined rheological and bond model shows a remarkable agreement with the experimental data for both rupture tension, strain, and time (**Fig. 7k - m**). Interestingly, simulations result in some variability (∼ 20% of the mean) that arises from the stochastic nature of the bond model. While the amplitude of this variability appears to depend on strain rate, it remains far smaller than the experimental variability (∼ 50-75% of the mean), indicating that biological variability also contributes.

The relationship between rupture tension and rupture strain (**Fig. 7h**) connects two qualitatively different domains. At high rates, the rupture tension is large and the rupture strain is relatively low and constant. This is a regime where intermediate filament bundles are massively recruited and cannot relax, forcefully breaking the bonds. At slow rates, the rupture tension is smaller but the material is able to achieve much larger deformations before failing due to significant dissipation of stress in the intermediate filament network that delays the onset of rupture.

### The strain stiffening threshold depends on strain history

One fundamental hypothesis of our model is that keratin filament bundles relax stress to give rise to a shear stiffening behaviour. In vitro, keratin bundles have been shown to dissipate stress through interfilament sliding [59]. One implication is that the strain stiffening threshold should depend upon the strain history when the keratin supracellular network is present.

To test this, we subjected monolayers to a ramp of deformation to 100% strain at 1% s^−1^ strain rate, maintained them stretched for 20 min, and returned them to their initial length. We then repeated this procedure increasing the deformation to 125% and then 150% in cycles 2 and 3 (**Fig. S13a**-(i)). If our rheological model is conceptually correct, we expect that the threshold strain for stiffening will increase in cycles 2 and 3 to a strain close to the holding strain in the previous cycle because some interfilament sliding has taken place (or equivalently because dashpots have had time to flow in our rheological model, **Fig. 7h**), effectively adding extra length to the slack. In our experiments, the monolayers displayed strain stiffening around 50% deformation in cycle 1 and their stress relaxed by ∼ 60% over a period of ∼ 10 min (**Fig. S13a**-(i)). Consistent with our hypothesis, strain stiffening occurred around 75% in cycle 2 and around 85% in cycle 3 (arrows, **Fig. S13a**-(ii)). Stress relaxation with similar characteristics to the first cycle could be observed while the monolayers were maintained stretched in cycles 2 and 3 (**Fig. S13a**-(i)). In contrast, no such behaviours were observed in monolayers expressing K14-R125C (**Fig. S13b**), confirming that it was linked to the presence of an intact keratin supracellular network. Overall, these experiments suggested that our rheological model was qualitatively correct.

### Keratin filaments and desmosomes may reinforce intercellular adhesion to protect monolayers against rupture

How the presence of a keratin supracellular network connected by desmosomes protects monolayers against rupture and why monolayers can rupture at low stress when they increase their contractility remains unclear. To examine these questions, we used our intercellular junction model to probe the strength of adhesion between cells in different experimental conditions. Our model junction is parameterised by three physical values: *k_off_* /*k_on_* the dissociation constant of unloaded bonds, *f*_0_ the force dependence of the slip bond behaviour, and *N* the number of bonds. All of these parameters influence the strength of the monolayer (**Fig. S14a**). As there is considerable homology between the ectodomains 1-2 of E-cadherin and desmosomal cadherins [60, 61], we assumed that the dissociation constant was the same for all intercellular bonds. As E-cadherins and desmosomal cadherins show very similar slip bond behaviours above a threshold force of 30 pN [62, 63], we chose *f*_0_ identical for all bonds. As a consequence, we only varied the number of bonds *N* when adjusting our model to fit experimental observations (see methods). For this, we subjected our junction model to experimentally measured stress as a function of time curves and adjusted *N* such that ruptures had a 50% probability of occurring before the experimentally observed rupture stress and 50% after, signifying that we had reached the correct adhesive strength. Note that this number provides a relative scale rather than an absolute number of bonds. Another benefit of this approach is to allow us to probe the strength of intercellular adhesion between cells in experiments without necessitating knowledge of tissue rheology, which is particularly useful for the calyculin experiments.

First, we applied our approach to determine if the strength of adhesion depends on strain rate in wild-type monolayers. We reasoned that, as the deformation is large in all experiments, intermediate filaments and desmosomes should contribute to intercellular adhesion at all strain rates. Our model revealed no significant differences in the number of bonds participating to adhesion (**Fig. S14b-g**). Next, we compared the strength of adhesion in wild-type monolayers and those expressing K14-R125C (in which the keratin supracellular network is disrupted) subjected to ramps of deformation at 1% s^−1^. The number of bonds *N* involved in intercellular adhesion in control monolayers was ∼three-fold larger than in monolayers expressing K14-R125C (**Fig. 7n**), suggesting that disruption of the keratin intermediate filament network may lead to a lower contribution of desmosomes to intercellular adhesion. Again, the number of bonds in monolayers expressing K14-R125C was independent of strain rate (**Fig.S14h-i, k**). Finally, we reasoned that intermediate filaments and desmosomes would not be solicited in wild-type monolayers treated with calyculin because of the absence of tissue-scale deformation. Therefore, they should possess an intercellular adhesion strength similar to K14-R125C monolayers. When we fitted these experiments with our model, we found that *N* was not significantly different from that found in K14-R125C monolayers subjected to deformation, but significantly lower than in wild-type monolayers deformed at 1% s^−1^ (**Fig. 7n**, **Fig. S14j**). To confirm this, we experimentally characterised the response of K14-R125C monolayers to calyculin treatment and found no significant differences in the rupture tension or rupture time compared to wild-type monolayers treated with calyculin (**Fig. S9h-j**).

These results suggest that the response of both K14 mutants and calyculin treated monolayers is not influenced much by IFs and desmosomes, and therefore dominated by actomyosin and cadherins. This implies that rheological models calibrated for actomyosin combined with the K14/calyculin bond model should also predict the correct distribution of strength and rupture strains; **Fig. 7o** shows that only a small adjustment of the parameters of the actomyosin driven rheological model (see inset **Fig. 7e**) is required to match model and experimental data.

Together these data suggest that the application of strain to wild-type monolayers engages a additional cytoskeletal network involving keratin intermediate filaments and increases the strength of intercellular adhesion due to the involvement of additional bonds, likely due to desmosomal cadherins.

## Discussion

We have characterised fracture in epithelial monolayers and showed that they are remarkably strong, withstanding several-fold increases in length before the initiation of cracking. Remarkably, increases in cell contractility also ruptured monolayers in the absence of any applied deformation. By systematically varying strain rate, we reveal a trade-off between rupture tension and rupture time with large tensions leading to short lifetimes. Importantly, we unravelled a role for the supracellular keratin filament network in controlling the rheology and strength of tissues both in vitro and in vivo. Finally, using computational modelling, we show that rupture onset depends strongly on tissue rheology and collective bond dynamics under force.

Our experiments revealed a key role for keratin intermediate filaments in the response of tissues to large deformations: they governed strain stiffening and protected monolayers against early rupture. Importantly, perturbing the keratin intermediate filaments directly or indirectly by disrupting their interfacing to desmosomes had the same qualitative effect, signifying that it is the supracellular network linking individual cellular keratin networks that is crucial for tissue strength and strain stiffening. Consistent with our in vitro experiments, combined depletion of keratin 8 and desmoplakin in Xenopus embryos fragilised the epidermis, leading to the formation of cracks during body axis elongation. Whereas the actomyosin cytoskeleton is central to cell and tissue mechanics at low strain [64, 65, 20], its perturbation did not affect strain stiffening nor the strength of tissues. Most epithelia are bound to a basement membrane and, as a consequence, rupture of the epithelium *in vivo* will likely be influenced by the mechanical properties of the basement membrane. Future work will be necessary to characterise the strength epithelium-basement membrane composite.

These data paint a picture in which the actomyosin cytoskeleton controls tissue rheology for deformations smaller than ∼ 50% and keratin intermediate filaments dominate for strains above 100%, consistent with previous work [38]. This transition was manifested by a progressive stiffening of monolayers for strain above 50%. Interestingly, such a mechanism was not observed in previous work examining the response of epithelia to large deformation [38], perhaps because of the much longer time-scales over which strain was applied (tens of hours) resulting in a very low strain rate. The exact mechanism through which strain stiffening arises remains to be determined. However, we hypothesise that it is due to progressive tensile loading of keratin bundles with increasing deformation, a mechanism commonly observed in random fibre networks [66, 67, 68] and proposed to play a role in tissues [69]. Previous work has shown that, at low strain, keratin intermediate filament bundles appear wavy but straighten when deformation increases, indicative of a transition from unloaded to loaded [38, 44]. Within cells, the keratin network radiates from the perinuclear region towards the cell junctions [70] (**Fig. 5a, e**). In response to uniaxial stretch, keratin filaments aligned with the direction of stretch will straighten first and, as deformation increases, filaments at increasingly larger angles from this direction will straighten. Thus, more and more mechanical elements bear load as deformation increases. During physiological function, strain stiffening may help epithelia limit how much they deform in response to an external force. Indeed, with strain stiffening, each additional increment in deformation necessitates the application of a larger increment in force.

Our experiments show that strain stiffening is strain rate dependent. This signifies that, at low strain rate, strain stiffening is absent and this may allow tissues to deform substantially. Experiments in which we subjected monolayers to cycles of increasingly large deformation suggested that the keratin network remodels over tens of minutes to dissipate stress (**Fig. S13**). The molecular mechanism underlying strain rate dependency remains unclear; however, it might involve molecular turnover of proteins within the keratin-desmosome force chain or sliding between filaments within the keratin bundles. Indeed, experiments examining the mechanical response of keratin bundles in cells and in vitro both indicate the presence of sliding between filament subunits in response to stretch [40, 59]. Alternatively, turnover of proteins within the cytoskeleton has been shown to dissipate stress. At low strain, cells and tissues dissipate stress on minute time-scales due to turnover of actomyosin [71, 20]. Monolayers subjected to 0.1% s^−1^ do not strain-stiffen, indicating that potential dissipatory turnover mechanisms act faster than the ∼ 10 - 20 minutes necessary to reach the strain magnitudes at which stiffening is observed. While keratins and desmoplakin turn over with characteristic times of ∼ 1h [72, 20], the desmosomal cadherin desmoglein 2 is reported to turn over on a time-scale of ∼ 20 minutes [73] and plakophilin turns over substantially within 5 minutes [74]. Future work will be necessary to determine which of interfilament sliding or molecular turnover contributes most to stress dissipation in keratin networks.

Together our experiments and modelling allowed to link rupture of bonds at the molecular-scale to cellular forces arising from tissue-scale deformation of the monolayer. While the trade-off between force and lifetime was expected from previous work [24], our experiments indicate that tissue rheology plays an integral part in defining rupture onset. Remarkably, our model shows that prediction of rupture tension and strain necessitates the implementation of a realistic tissue rheology that incorporates shear stiffening. Our data poses an intriguing question. At high strain imposed at sufficiently high strain rate, wild-type monolayers strain stiffen and, as a result, their tension is larger than monolayers with a perturbed keratin network. If the number of adhesion proteins linking the cells is the same in both conditions, the force that each bond must bear will be larger in wild-type tissues than in K14-R125C monolayers. As a consequence, we would expect wild-type tissues to rupture at lower strains than keratin compromised ones, the contrary of what we observe. One potential explanation may be that desmosomal cadherins or desmosomal proteins possess catch-bond properties, similar to E-cadherin and alpha-catenin [62, 75]. At zero force, catch bonds have a very short lifetime but, as applied force increases, their lifetime grows to an optimum before decreasing again. Work using FRET tension sensors has shown that, under resting conditions in adherent monolayers, adherens junctions are under tension but that desmosomes are not [34, 42]. Thus, at low strain when keratin filaments are unloaded, desmosomal cadherins may not sense any force and have a lifetime so short that their contribution to intercellular adhesion is minimal. As strain increases, keratin filaments become progressively loaded, exerting tension on desmosomes [42] and potentially increasing the lifetime of desmosomal cadherins as well as their contribution to load bearing. Thus, stretch would lead to both strain stiffening and an increase in the effective intercellular adhesion. Consistent with this hypothesis, our computational model of intercellular junctions predicts that rupture stress decreases with decreasing number of bonds (**Fig. 14a**) and that wild-type monolayers have more intercellular adhesive bonds than monolayers with disrupted keratin networks (**Fig. 7n**). Furthermore, transcriptomic data indicates that the number of transcripts for E-cadherin, desmoglein 2, and desmocollin 2 are comparable in MDCK cells, signifying that desmosomal cadherins could potentially provide these extra bonds (**Table S6**). Conversely, when stress arises from an increase in myosin contractility rather than deformation, no additional bond recruitment takes place in wild-type monolayers (**Fig. 7n**). Together these data suggest that the effective number of intercellular adhesion bonds increases when deformation is applied to monolayers in which a supracellular keratin network is present. However, we note that other changes to the model parameters (such as the slip bond characteristic force *f*_0_ or the dissociation constant) can also lead to similar changes in rupture characteristics (**Fig. 14a**). Future work will be needed to thoroughly investigate the mechanism of adhesive strength reinforcement and experimentally characterise the associated physical and biological parameters.

From a physiological point of view, the shear-stiffening behaviour enables the tissue to respond very differently to mechanical perturbations depending on the strain rate (**Fig. 7p**). The tissue responds to a fast, shock-like, perturbation by stiffening and increasing effective adhesive strength, therefore limiting the deformation and maximising the force at which the material fails. However, when subjected to a slow and steady deformation, the material can tolerate very large stretch without failure. This is, to our knowledge, the first characterisation of such a dynamic transition with regards to rupture behaviour but the exact biophysical mechanisms underlying it remain unclear. Our computational model assumes that the number of intercellular adhesive proteins within a junction does not change during deformation. Yet, imaging reveals that the height of intercellular junctions visibly decreases with strain (**Fig. 1f**). How changes in junction shape and size affect the number of E-cadherins and desmosomal cadherins engaged in intercellular adhesion as well as their stability is unclear. Previous work examining shrinkage of intercellular junctions in response to myosin contractility have reported an increase in the density of E-cadherins, suggesting that the overall number of E-cadherin links present at the junction remains constant [76]. In addition, application of stress can modulate the stability of intercellular junctions by decreasing cadherin turnover [77], increasing the life time of proteins within the cadherin-catenin complexes (e.g. E-cadherin and alpha-catenin, [62, 75, 78]), and recruiting proteins to reinforce intercellular junctions (e.g. vinculin) [79]. Finally, our study further suggests that application of deformation increases the effective number of intercellular bonds by putting the keratin intermediate filament network and desmosomes under mechanical load. Thus, in principle, deformation could lead to changes in the number of intercellular bonds *N*, their dissociation constant, and their sensitivity to force *f*_0_. Further work will therefore be necessary to determine the exact contribution of each of these mechanisms to the overall strength of tissues and its modulation by strain application.

In summary, our work has shown that the mechanics of tissues at high strain and high strain rate is dominated by a supracellular network of keratin intermediate filaments linked by desmosomes. This network protects monolayers from rupture by limiting deformation through strain stiffening and may also increase effective intercellular adhesion when it is mechanically loaded. One implication of our work is that the rupture characteristics of monolayers expressing K14-R125C reflect the strength of actomyosin and adherens junctions, while the rupture characteristics of monolayers treated with latrunculin reflect the strength of keratins and desmosomes. In the present study, we only examined the onset of rupture and future work will be needed to investigate crack propagation in the plane of the tissue.

## Acknowledgements

The authors thank past and present members of the Charras and Kabla labs for discussions. The authors acknowledge technical support in LabVIEW development from A. Lisica. JD and GC were supported by a European Research Council consolidator grant (CoG-647186). JD and LB were supported by a sLOLA grant from the British Biotechnology and Biological Sciences Research council (BBSRC, BB/V019015/1) to GC. AB was supported by the seal of Excellence (SoE) fellowship from Politecnico di Milano. JF was supported by a grant from the BBSRC (BB/M003280 and BB/M002578) to GC and AK. AH was supported by a BBSRC grant (BB/K013521) to GC and AK. EF was supported by BBSRC grant (BB/V019015/1) to GC. EHB and SRA were supported by European Research Council Starting Grant (ERC-StG) under the European Union’s Horizon 2020 research and innovation programme, Grant agreement No. 950254 (to EHB); The European Molecular Biology Organization (EMBO) Installation Grant, Project No. 4765 (to EHB); La Caixa Junior Leader Incoming, No. 94978 (to EHB); Instituto Gulbenkian de Cîencia (IGC) and Fundação Calouste Gulbenkian (FCG), start-up grant I-411133.01 (to EHB); and F FCT PhD Fellowship UI/BD/152259/2021 (to S.R.A.). IGC’s Advanced imaging (PPBI-POCI-01-0145-FEDER-022122), Genomics (LISBOA-01-0246-FEDER-000037), Bioinformatics and Aquatic animal facilities. EHB was partially funded by the Deutsche Forschungsgemeinschaft (DFG, German Research Foundation) under Germany’s Excellence Strategy (EXC 2068, 390729961, Cluster of Excellence Physics of Life of TU Dresden).

## Author Contributions

JD, JF, AK and GC conceived the project and wrote the manuscript. JD performed most of the experiments, analysis, and microscopy. JF performed some experiments and provided experimental and analysis tools. AH established the initial methods and contributed some experiments. EF performed the Western blotting and helped with establishment of cell lines. LB performed experiments examining the response of K14 R125C monolayers to calyculin. EHB and SRA performed experiments in Xenopus. GC contributed reagents and established some of the cell lines. AB and AK designed and implemented the model. AK advised on analysis. AK and GC oversaw the entire project and writing. All authors discussed the results and manuscript.

## Data availability

All reagents are available from the corresponding author upon request. Code and primary data are available from the UCL data repository (https://rdr.ucl.ac.uk/).

## Methods

### Cell culture

MDCK cells were cultured at 37 °C in an atmosphere of 5% CO_2_ in DMEM (1X) + Glutamax (Thermo Fisher) supplemented with 10% fetal bovine serum (FBS, Sigma-Aldrich), 2.5% of 1M HEPES buffer (Sigma-Aldrich) and 1% penicillin-streptomycin (Thermo Fisher). Cells were passaged at 1:5 ratio every 4 days using standard cell culture protocols and disposed of after 30 passages. Mechanical experiments were performed in Leibovitz’s L15 without phenol red (Thermo Fisher) supplemented with 10% fetal bovine serum, 2.5% of 1M HEPES buffer and 1% penicillin-streptomycin. For imaging and mechanical testing, the culture medium was exchanged for imaging medium that consisted of Leibovitz L15 without phenol red supplemented with 10% FBS.

To visualise junctional and cytoskeletal structures, we used stables lines expressing E-cadherin-GFP, Vinculin-GFP, EPLIN-GFP, and Keratin 18-GFP. Details about their generation are given in [20], [44]. Stable expression of the tagged protein of interest was ensured by antibiotic selection using either 250 ng/mL puromycin or 1 mg/mL G418.

To study the role of intermediate filaments, we generated cell lines stably expressing Keratin 14-R125C, which leads to disruption of the keratin network. The cDNA encoding Keratin 14 R125C-YFP was a kind gift of Thomas Magin (University of Leipzig, Germany) and was cloned into a retroviral vector (pTRE, Takara Clontech). Retrovirus was generated as described in [44] and transduced into MDCK cells. After 2 weeks selection with hygromycin (400 µg/mL), cells were sorted to achieve a homogenous level of fluorescence.

To deplete desmoplakin, we purchased shRNAs targeting dog desmoplakin in a lentiviral vector (V3LHS 302846 and 302847, pGIPZ vector, Horizon discovery, Perkin Elmer, UK). We generated lentiviral particles following manufacturer instructions, transduced MDCK cells, selected cells with puromycin and sorted the cells with flow cytometry to achieve a homogenous knock down. As a control, we also generated a cell line expressing non-silencing shRNA using the same methods. Protein depletion was verified by immunoblotting (**Fig. S7**). The antibodies used were anti-desmoplakin (Progen) and with anti-GAPDH as a loading control (Abcam). Appropriate HRP-coupled secondary antibodies (dilution 1:10,000) were from Jackson ImmunoResearch and Cytiva. The shRNA used in this work was V3LHS302847, shRNA2 on **Fig. S7**.

None of the cell lines in this study were found in the database of commonly misidentified cell lines maintained by International Cell Line Authentication Committee and National Center for Biotechnology Information Biosample.

All lines were routinely screened for the presence of mycoplasma using the MycoALERT kit (Lonza).

### RNA sequencing of MDCK cells

We used mRNA sequencing (RNASeq) to quantify the normalised expression of mRNA transcripts for proteins in subcellular structures as described in [20]. Briefly, to prepare total RNA samples,

MDCK cells were cultured for 3 days to reach confluence. This provides sufficient time for the junctions to mature and for the mRNA content to be regulated. Next, the total RNA was extracted using Tri reagent (Sigma Aldrich) following the manufacturer’s protocol. Samples were processed using Illumin’s TruSeq Stranded mRNA LT sample preparation kit (p/n RS-122-2101) according to manufacturer’s instrucitons. Samples were sequenced on the NextSeq 500 instrument (Illumina, San Diego, USA) using a 43bp paired end run resulting in ¿15 million reads per sample. resulting in ¿15 million reads per sample. Run data were demultiplexed and converted to fastq files using Illumina’s bcl2fastq Conversion Software v2.16. Fastq files were then aligned to the CanFam3.1 assembly released by the Dog Genome Sequencing Consortium using Tophat 2.014 then deduplicated using Picard Tools 1.79. Reads per transcript were counted using HTSeq and normalised expression for each mRNA transcript was estimated using the BioConductor package DESeq2.

### Stress measurement devices

The stress measurement devices were an adaptation of the force measurement device described in [19]. Briefly, two nickel-titanium (nitinol) wires (Euroflex) with different stiffnesses were glued into a bent glass capillary (Sutter Instruments). The arm with the stiffer wire was covered by a glass capillary to create a reference rod and two Tygon cylinders were glued to the extremities of both wires.

### Generation of suspended epithelial monolayers

Suspended epithelial monolayers were generated as described by [19]. Briefly, mechanical devices were glued into 50 mm diameter Petri dishes, placing a glass capillary underneath them to prevent contact between the device and the bottom of the Petri dish which creates friction. To maintain the distance between the rods constant during the preparation procedure, a custom-designed, 3D printed plastic holder was placed in between them.

Collagen was reconstituted on ice in the following v/v proportions: 50% collagen (Cellmatrix type I-A, Nitta Gelatin), 20% 5X DMEM, 20% sterile water, and 10% reconstitution buffer (50 mM NaOH solution in sterile water, 200 mM HEPES and 262 mM of NaHCO_3_) following manufacturer instructions. A 10 µL drop of collagen was placed between the rods and left to solidify in a dry incubator at 37 °C for 1-1.5 h. Once a solid collagen scaffold was formed, it was rehydrated by placing an 8 µL drop of cell culture medium onto it and two 250 µL drops in the bottom of the dish. The dish was then placed for 30 min inside a humidified incubator at 37 °C. During the rehydration time, confluent flasks of MDCK cells were trypsinised for 20 min. Cells were then resuspended to a final concentration of 3×10^4^ cells per 10 µL. After rehydration, a 10 µL drop of the resuspended cells was placed on top of the collagen scaffold; cells were left to settle onto the collagen for 30 min inside the incubator. After this time, 8 mL of medium was added to each Petri dish, and both the V-shaped glass capillary and the holder separator were gently removed. The devices were left in the incubator for 48 - 72 h to allow cells to grow to confluence, covering the collagen scaffold and part of the Tygon cylinders on each test rod.

### Removal of the collagen substrate

Immediately prior to experimentation, a collagenase solution was prepared by mixing collagenase type-II (Worthington Biochemical) with imaging medium to reach a final concentration of 250 units per mL. This solution was gradually exchanged with the cell culture medium in the Petri dishes containing the devices and then left for 1 h at 37 °C to allow for full enzymatic digestion of the collagen. Finally, the collagenase solution was gradually replaced with imaging medium. The device was then ready to be used for experiments.

### Mechanical testing procedure

The mechanical setup was mounted on the stage of an inverted microscope (Olympus IX-71) and is described in [20]. The stiffer rod of the device was brought into contact with the arm of a motorized manipulator (M126-DG1) controlled through a C-863 controller (Physik Instrumente) whereas the softer arm of the device was attached to the tip of the force transducer (SI-KG7A, World Precision Instruments) held in position by a manual micromanipulator. Both the motorized manipulator and the force transducer were mounted onto magnetic plates to secure them firmly onto the microscope stage. The motorized manipulator allowed us to subject monolayers to different strains with precise control of strain rate. Stretched monolayers exerted restoring forces on the flexible rod, deflecting the force transducer. This deflection was transformed into a voltage that was converted into a digital signal using a data acquisition system (USB-1608G, Measurement Computing) and recorded on a computer. The motorized manipulator was controlled using a custom-written code in Labview (National Instruments). During the experiment, images of the monolayer were taken every 1 s using a 2X objective (2X PLN Olympus) and a GS3-U3-60QS6M-C, Point Grey camera.

### Quantification of tissue tension

In experiments using suspended epithelial monolayers, the output force measured by the transducer is in Volts and we converted it into force in Newtons. To do this, each experiment was individually calibrated. After each experiment, monolayers were broken if they had not already failed. In these conditions, all of the force measured by the force transducer is due to the deflection of the soft wire, *d_W_*, and it can be determined using the force-deflection equation for a simple cantilever beam.

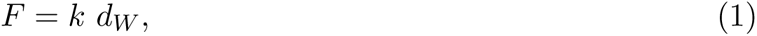

where *k* is the stiffness of the wire defined as:

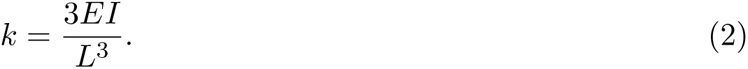

*E* is the elastic modulus of the wire (previously determined in [19]), *I* its moment of inertia and *L* its length.

To determine the conversion between Volts and Newtons, for each experiment we collected 6 voltage, deflection pairs (*V*, *d_W_*) and fitted them to a linear function. Using this procedure, we could determine the conversion factor *α* to convert Volts into Newtons:

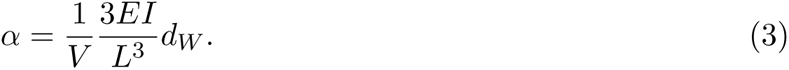

In our mechanical characterisation of monolayers, we decided to approximate the tissue to a thin two-dimensional sheet and normalised the force *F* exerted on the monolayer to the average width of the monolayer before stretch *w*_0_,

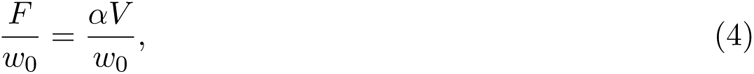

where *w*_0_ was sampled from three positions in the monolayers and computed as

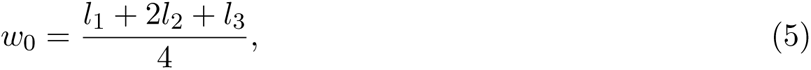

where *l*_1_ and *l*_3_ correspond to the width of the monolayer on each of the sides where it contacts the rods and *l*_2_ is the width at the middle point of the monolayer (**Fig. S1** d, inset). This definition of width was chosen because the location of the first rupture was unpredictable and did not always coincide with where the width was minimal.

All the tension measurements in this manuscript have been calculated using the initial width of the monolayer *w*_0_ as defined in eq.5 unless otherwise specified. Generally, tension measurements were smoothed with a moving average sliding fixed-time window around the time point of interest (30 points for strain rates above 1% s^−1^, 100 for 0.3% s^−1^ and 300 for 0.1% s^−1^).

Strain energy measurements were determined by integrating the area of the tension-strain curves up to the rupture point.

All the analysis was implemented in MATLAB R2019a (MathWorks, Natick, Ma, USA).

### Pre-tension measurements

Epithelial monolayers are intrinsically under tension due to forces generated by myosin contractility in the cells [20], [22]. These intrinsic forces generate a deflection on the flexible arm of the device that corresponds to the pre-tension of the monolayer Γ_0_. This pre-tension was determined from two bright-field microscopy images, one acquired at the beginning of the experiment and the other at the end once the monolayer was broken. Both images were taken when no parts of the mechanical testing system contacted any of the rods of the device. A stack of these two images was generated and a region of 250 × 150 pxl^2^ was cropped around the flexible arm to measure its displacement, Δ*x*, using a custom-written script in MATLAB. Using Hooke’s law the pre-tension of the monolayers is:

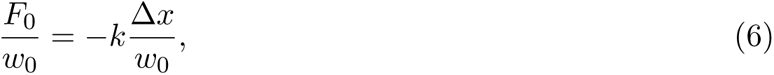

where *k* is given by eq.2.

### Tangent modulus measurements

To determine the tangent modulus of monolayers, we first fitted a smoothing spline to the experimental tension-strain curves. We then computed the derivative of this curve before fitting it with a smoothing spline to reduce noise. These operations were implemented in MATLAB R2019a, MathWorks.

### High magnification imaging devices

Devices used for confocal imaging to determine protein localisation as well as cell shape changes were similar to those described in [23]. Briefly, a glass capillary was bent into a U-shape using a small blow torch. One of the arms of the U-shaped capillary was cut at ∼ 5 mm from its base; in this arm, a nickel-titanium (nitinol) wire (Euroflex) was inserted to act as a hinge and covered by another piece of a glass capillary. Glass coverslips (VWR) were affixed using UV-curing glue (Loctite Glassbond, 447 Henkel) to the extremities of the glass capillaries to act as a substrate for cells to grow on. For precise control when stretching the monolayers, another piece of glass capillary was glued onto the end of the flexible arm at an angle to allow continuous contact with the manipulator arm (**Fig. 8**).

**Figure 8:**
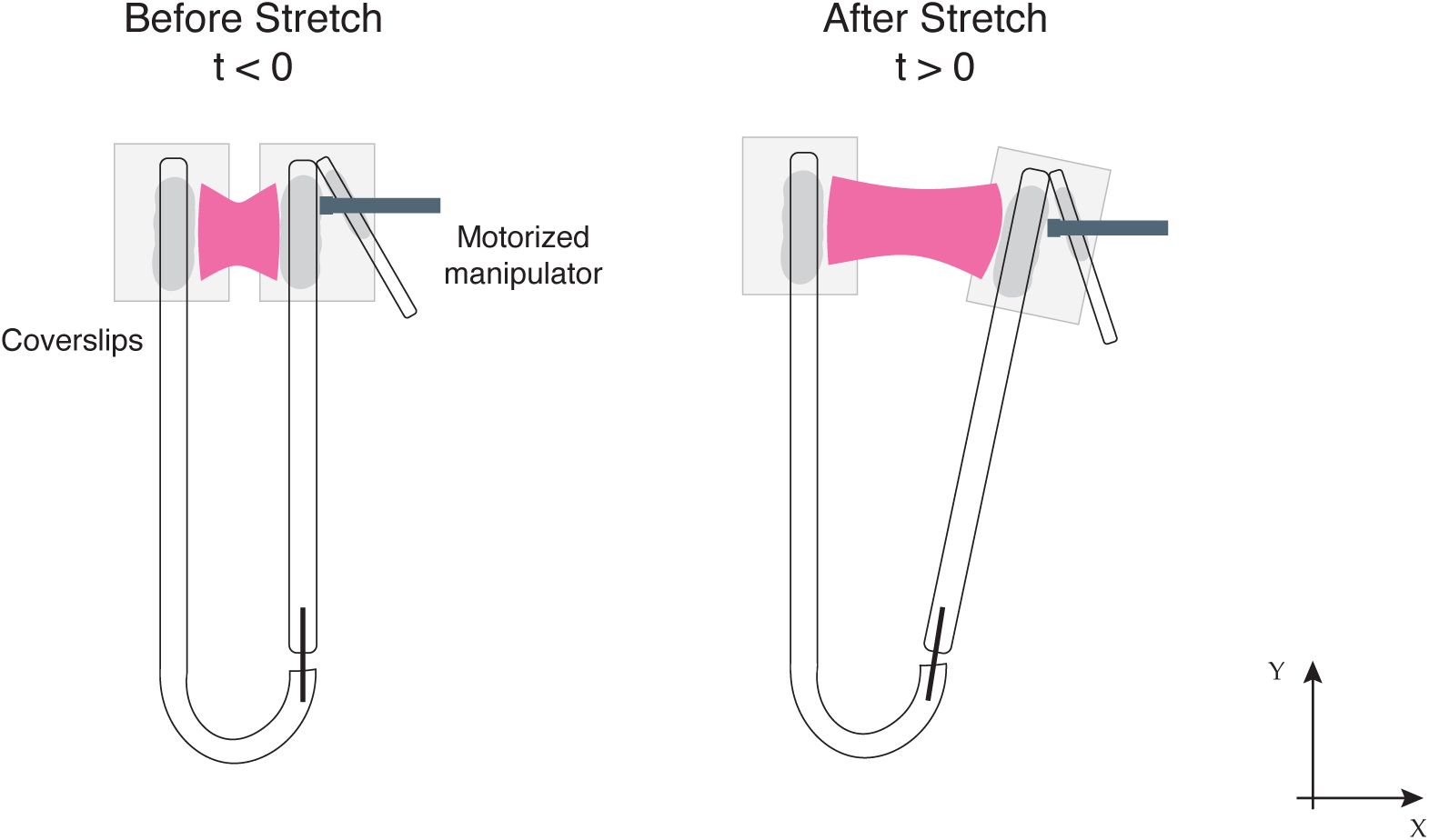
Example of a high magnification imaging stretching device. Shaded areas in dark gray depict the UV-glue used to affix the glass capillaries to the coverslips.

### Confocal imaging of tissues and mechanical manipulation

Epithelial monolayers were imaged at room temperature in imaging medium. To visualize cell membranes, tissues were incubated for 10 min with CellMask orange membrane stain following the manufacturer protocol (Thermo Fisher Scientific). To visualize cell-shape changes by dye exclusion, AlexaFluor-647-conjugated dextran, 10,000 MW (Thermo Fisher Scientific) was added to the imaging medium to a final concentration of 20 µg mL^−1^. Confocal images were acquired using a 60X Sil objective (UPLSAPO-S, numerical aperture 1.3, Olympus) mounted on an Olympus IX83 inverted microscope equipped with a scanning laser confocal head (Olympus FV-1200). Images consisted of a z-stack acquired at a spatial interval of 0.5 - 1µm. To generate time series, stacks were acquired every 50 s during stretching experiments and every 5 min in experiments in which monolayers were treated with calyculin. High magnification bright field images in **Fig. 1g** were taken using a 40X objective (Olympus, LUCPlanFL N) on an inverted Olympus IX71 microscope equipped with a GS3-U3-60QS6M-C, Point Grey camera.

### Immunohistochemistry assays

To visualise the organisation of the cytoskeleton and junctional proteins, we used immunostaining. For imaging of intermediate filaments, cells were fixed with a 1:1 mix of methanol and acetone at −20 °C for 10 minutes. For all other proteins, cells were fixed in 4% paraformaldehyde diluted in DMEM without phenol red for 15 minutes. Cells were then washed three times with PBS to remove any fixative. Cells were permeabilised with 0.5% Triton 100X in PBS for 5 min at room temperature. After permeabilisation, cells were washed three times with PBS and blocked with 10% horse serum (HS) in PBS for 1h at room temperature changing the blocking buffer every 15 min. Next, cells were incubated with the primary antibodies for 2h at room temperature in a solution of 10% HS in PBS. After washing three times with PBS, the cells were incubated with phalloidin 647 or 568 (Life technologies, 1:500) along with the appropriate alexa-conjugated secondary antibodies for 1h at room temperature in a solution of 10% HS in PBS. Finally, cells were washed three times with PBS. The following primary antibodies were used: mouse anti-phospho-myosin light chain 2 (S19) (Cell Signaling 3675S, 1:100 dilution), mouse anti-E-cadherin (BD Biosciences 610181, 1:200 dilution), mouse anti-cytokeratin-18 (abcam ab668, 1:100 dilution), rabbit anti-alpha-catenin (Sigma Aldrich C-2081, 1:200), and rabbit anti-desmoplakin (abcam ab71690, 1:100).

### Drug treatments

To block myosin contractility, blebbistatin (Sigma-Aldrich) was added at 50 µM concentration. To increase myosin contractility, we inhibited phosphatases using calyculin A (Sigma-Aldrich) at 20 nM. To block actin polymerization, latrunculin B (Calbiochem) was used at 1 µM. DMSO was added to control monolayers accordingly. Drug treatments were started 20 minutes before performing the ramps in deformation. To study ruptures caused by increases in internal contractility, calyculin was added at time 0 until full rupture of the monolayer. In immunostainings, drugs were added 15 minutes before fixation.

### Measurement of junctional protein recruitment in response to strain

#### Fluorescence Intensity Measurements in XYZ-t images

Z-stacks were acquired at 1 minute intervals on a confocal microscope, starting 5 minutes before stretch, and continuing for 30 - 80 minutes. The planes in which we measured the fluorescence intensity were selected by comparing a maximum intensity projection image (MIP) with all the planes at each time-point. This was done using 2D cross-correlation. Alignment between the last time point before stretch and the first one after stretch was done using 2D cross-correlation to ensure similar fields of view were compared. After this, a movie with the optimum z-planes is created and saved for further processing. Segmentation of cell membranes was carried out with Packing Analyzer to generate a mask containing all of the cell junctions. These masks were then processed in MATLAB. A second registration is performed to ensure the masks overlap correctly with each of the images of the movie. Finally, the intensity of each pixel within the mask was extracted from the aligned movies and averaged to output the mean fluorescence in the region of interest at each time point. All processing after segmentation was performed in MATLAB (as shown in **Fig. 9**)

**Figure 9:**
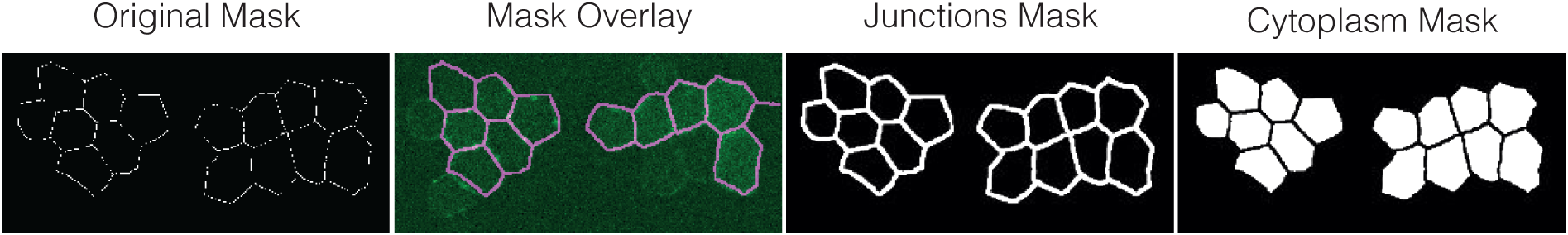
Processing of segmentation masks to perform fluorescence intensity measurements in either the junctions or the cytoplasm.

### Frog manipulation, embryo generation, and maintenance

Animal procedures were approved by the Ethics Committee and Animal Welfare Body (ORBEA) of the Instituto Gulbenkian de Cîencia (IGC), Portugal, and complied with the Portuguese (Decreto-Lei n° 113/2013) and European (Directive 2010/63/EU) legislations. *Xenopus laevis* oocytes were collected by inducing superovulation of mature females with human chorionic gonadotropin (Chorulon) [80]. Briefly, oocytes were fertilized using a sperm solution in Marc’s modified Ringer 0.1x medium (MMR: 10 mM NaCl, 0.2 mM CaCl2·2H2O, 0.2 mM KCl, 0.1 mM MgCl2·6H2O, and 0.5 mM HEPES, pH 7.1–7.2). After de-jellying, embryos were kept in 0.1x MMR at 12–21°C. The developmental stage of the embryos was constantly monitored and defined by following established developmental tables [81].

### Microinjection of frog embryos

Embryos were transferred into 5% ficoll (Sigma, P7798)/0.45x MMR (w/v)) prior to injection and morpholinos or mRNA were injected in the dorsal and ventral blastomeres at the 4-cell stage. All microinjections were performed using calibrated glass needles mounted onto a cell microinjector (MDI, PM1000) programmed to deliver 10 nL in a pulse of 0.2 seconds. To visualize nuclei and membranes of epithelial cells in vivo, 250pg of nuclear RFP and membrane GFP mRNA were injected per blastomere. Furthermore, to knockdown keratin 8 and desmoplakin, previously validated morpholinos (MO) [53, 54] were coinjected with membrane and nuclear markers. krt8-MO and dsp-MO were injected at a concentration of 300 µM per blastomere.

### Epidermis RNA library preparation and analysis

Epidermis of *Xenopus laevis* was isolated and processed for RNA extraction. Briefly, RNA quality was assessed in a HS RNA Screen Tape Analysis (Agilent Technologies), and mRNA-libraries were prepared using SMART-Seq2 kits. Illumina libraries were generated with the Nextera standard protocol. Libraries quality was assessed in a Fragment Analyzer (AATI). Sequencing was carried out in NextSeq500 Sequencer (Illumina) using 75 SE high throughput kit. Sequences were extracted in FastQ format using the bcl2fastq v2.19.1.403 (Illumina). After filtering for ribosomal contamination, sequences were mapped against the reference genome of Xenopus laevis XENLA-9.2-Xenbase.gtf (v9.2) (https://ftp.xenbase.org/pub/Genomics/JGI/Xenla9.2/). Gene expression tables were imported into the R v3.6.3 to normalize gene expression with the TMM (Trimmed Mean of M-values) procedure [82, 83] by using the NOISeq R package (v2.30.0) [84].

### Frog immunofluorescence

Embryos were fixed with Dent’s Fixative (20% DMSO, 80% Methanol) for 2 hours at room temperature with gentle agitation. After fixation, embryos were permeabilised with 1x PBS 0.3% Triton X-100 (v/v) for 30 minutes and blocked with 10% normal goat serum (NGS) in 1x PBS for 30 minutes at room temperature. Embryos were then incubated overnight at 4°C with 1:50 primary antibody (keratin type II, 1h5, Developmental Studies Hybridoma Bank). Embryos were washed with 1x PBS 0.3% Tween-20 3 times and incubated with secondary antibody (anti-mouse Alexa Fluor 647) at 1:350 and DAPI solution (62249, Thermo) at 1:1000 for 2 hours at room temperature. Embryos were then washed 3 times and fixed with PBS 4% Formaldehyde for 10 minutes at room temperature before imaging in a confocal microscope (described below).

### Frog embryo mounting, microscopy, and time-lapse imaging

Embryo mounting. Embryos were mounted and imaged in agarose wells. Wells were shaped using OD-1.5mm borosilicate glass capillaries in solidifying 1% Agar in 0.1x MMR. After solidification of the agar, the capillaries where carefully removed, and wells were filled with 3% Methyl Cellulose solution (3% methylcellulose in MMR 0.1x). Plates where then filled with MMR 0.1x and embryos were placed with the anterior part (head) pointing towards the end of the well. Fixed embryos were mounted in similar wells but filled with PBS.

Embryo extension live imaging. Z-stacks of live embryos were acquired on a Leica, Stellaris 5 up-right system using either a HC APO L U-V-I 10x/0.30 NA WATER (Leica) or a HC FLUOTAR L VISIR 25X/0.95 NA WATER (Leica) objective and the DPSS 561 and OPSL 488 lasers. Confocal image stacks of the embryos were acquired for 5 hours with intervals of 7.5 minutes. The system was controlled by LAS X (Leica).

Immunofluorescence and fixed embryo imaging. Z-stacks of live and fixed embryos were acquired on a Leica, Stellaris 5 upright system using a HC FLUOTAR L VISIR 25X/0.95 NA WATER (Leica) or a HC APO L U-V-I 40x/0.80 NA WATER (Leica) objective and the Diode 405, Diode 638, DPSS 561 and OPSL 488 lasers. The system was controlled by LAS X (Leica). Digital zoom was useed in some cases.

### Frog image processing and data processing

Image treatment and processing. Image level adjustment, morphological segmentation, stack projection and time-lapse videos, where performed using Fiji ImageJ built-in plugins. Photoshop and Illustrator were used to generate final figures.

Aspect ratio calculation. Aspect ratio was calculated using frames of early and late control or MO-injected embryos (imaged as described above). For all above mentioned conditions, the GFP tagged membranes of embryos constituted the input image and each individual cell was segmented using the Morphological Segmentation plugin in Fiji. Cells not automatically recognized by the segmentation plugin where manually segmented using the ROI manager. After proper segmentation, aspect ratios were accessed for each cell through the minimum bounding rectangle method. Cell strain calculation. Maximum strain was assessed at each time point in elongating live embryos by measuring the change in dimension of the cells along the anterior-posterior axis (deformation axis) using the formula: strain(t)=(Xt-Xi)/Xi with Xt the cellular AP length at time t and Xi the initial cellular AP length. All lengths were obtained using Fiji for each time point.

### Statistical Analysis

#### Suspended monolayers

Statistical analyses were performed using MATLAB R2019a (Mathworks, Natick, MA, USA). Box plots show the median of the distributions with a central bar, the 25th (first quartile, Q1) and 75th percentiles (third quartile, Q3) are represented by the bounding boxes and the most extreme data points without the outliers are represented by the whiskers. Outliers were defined as being either larger than *Q*3 + 1.5*IQR* or smaller than *Q*1 − 1.5*IQR*, with *IQR* = *Q*3 − *Q*1. They appear outside the range of the whiskers and are represented by the symbol ‘+’ in red. In all box plots, statistically significant difference are marked as one of: ns non significant, P *>* 0.05, *P *<* 0.05, **P *<* 0.01, ***P *<* 0.001, ****P *<* 0.0001. Statistical significances were computed using a Kolmogorov-Smirnov test. The number of monolayers (unless otherwise specified in the legend) examined in each condition is indicated above each box plot.

#### Frog embryos

Data was represented and tested for normality and significance using Prism10 (GraphPad). Data sets were tested for normality using the d’Agostino–Pearson and/or Shapiro–Wilk test. When the distributions followed a normal distribution, significance was accessed using a students t-test (Two-tailed, unequal variances). When they did not, significance was calculated using a Kolmogorov-Smirnov-test (Two-tailed, unequal variances).

### Computational model

#### Molecular bond dynamics model

The rupture of a junction is modelled from the dynamics of a population of *N* links, each having a slip bond behaviour, i.e. a rate of binding *k_on_* that is assumed to be constant, and a rate of unbinding *k_off_* that is force dependent, *k_off_* = *k_off,_*_0_ exp (*f/f*_0_), where *f* is the force on the link [85]. In this implementation, the force on the junction *F* is distributed evenly over all of the bound links. If, at a certain time, there is a number *n_b_* of bound links, the force per link is *f* = *F/n_b_*.

The system is first initialised with a random distribution of states across the *N* links, with a probability to be bound given by *k_on_/k_off,_*_0_. The overall force over time *F* (*t*) follows precalculated curves based on the selected rheology or on experimental data. It is here assumed that stress is homogeneously distributed in all intercellular junctions, and therefore that the junction force *F* is simply proportional to the tissue tension Γ.

To simulate the evolution of the number of bound links within the population, a small time step *dt* and discrete probabilities that junctions change state are defined at each time point based on the current force. The state of the system then evolves in a stochastic manner based on these probabilities. The model runs until the end time point is reached, or until all links are unbound, which corresponds to the rupture time *t*^∗^. The model has four parameters. We fixed two parameters in the bulk of the analysis, the number of linkers *N* = *N_ref_* = 100, and set the ratio *k_on_/k_off,_*_0_ = 10. The values of the force *f*_0_ and *k_on_* were then varied to best match the experimental data for Γ^∗^ and *ε*^∗^ as a function of strain rate, once combined with the non-linear visco-elastic model presented below. The parameter values are presented in **Table S7**.

The model parameters predicted above do not capture the variability from one curve to the next. In order to estimate this variability, we considered individual experiments, and looked for the value of *N* that would best predict their rupture tension. In this situation, we only look at the data for Γ(*t*), and change *N* until the model predicts that 50% of the simulations fail before the experiment, i.e. we look for the value of *N* such that the experimental data is the median of the model distribution. We have to use such a criterion because if the model does not fail when Γ^∗^ is reached, there is no experimental data to extrapolate the behaviour. This approach was validated by demonstrating that the values of *N* obtained on all the wild type curves were distributed about our reference value *N_ref_* = 100, with no systematic trend as the rate is varied (**Fig. S14b-g**). When the same approach is deployed on perturbed monolayers (expressing K14-R125C or treated with calyculin), we find however that a significantly lower value for *N* is predicted in all these cases (**Fig. S14h-k**).

#### Linear and non-linear rheology

In the experiments and the model, the strain is controlled, and the mechanical tension Γ(*t*) of the epithelium is calculated using a rheological model. For a given strain rate *ε̈*, the strain function is given by *ε*(*t <* 0) = 0 and *ε*(*t* ≥ 0) = *ε̈t*. For the linear spring model, the tension is proportional to the strain: Γ = *k_a_ε*. For the fractional model, the calculation of the tension follows the method presented in [55], calculated using the software package RHEOS [86].

The non-linear spring model corresponds to a monotonically increasing relationship between Γ and *ε*. To define this relationship, we assume that strain-stiffening arises from the progressive recruitment of intermediate filaments as the strain is increased (in addition to the linear term defined in the previous paragraph). The strains at which fibers are recruited are modelled as a compact triangular distribution *p_s_*(*ε*) over a domain [0, 2.2], with a maximum in the middle (**Fig. S12d**). These bounds are based on the range of strains for which strain stiffening is observed in the experiments at high strain rates. The tension in the tissue is then calculated by superposition:

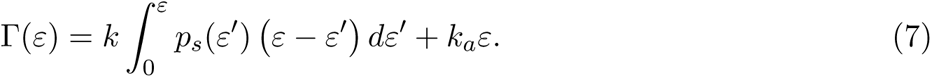

The non-linear viscoelastic model builds on the previous description, but assuming that once loaded, each filament behaves in a Maxwell-like manner. For a single Maxwell model with a spring *k* and a dashpot *η*, the response to the ramp in strain with rate *ε̈* is given by *ηε̈*(1 − exp(−*t/τ*)) with *τ* = *η/k*. The full response is again obtained by superposition:

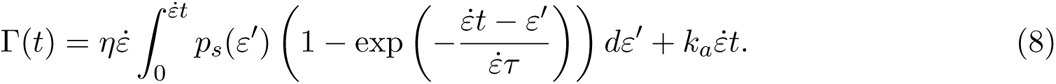

The distribution *p_s_* is adjusted to mimic the high strain rate limit, identical to the non-linear spring model (**Fig. S12d**). The extra parameter *η* (or equivalently *τ*) is adjusted to account for the observed timescale associated with the shear stiffening behaviour. The parameter values used in the different rheological models are presented in table **Table S8**.

## Supplementary Material

**Figure S1:**
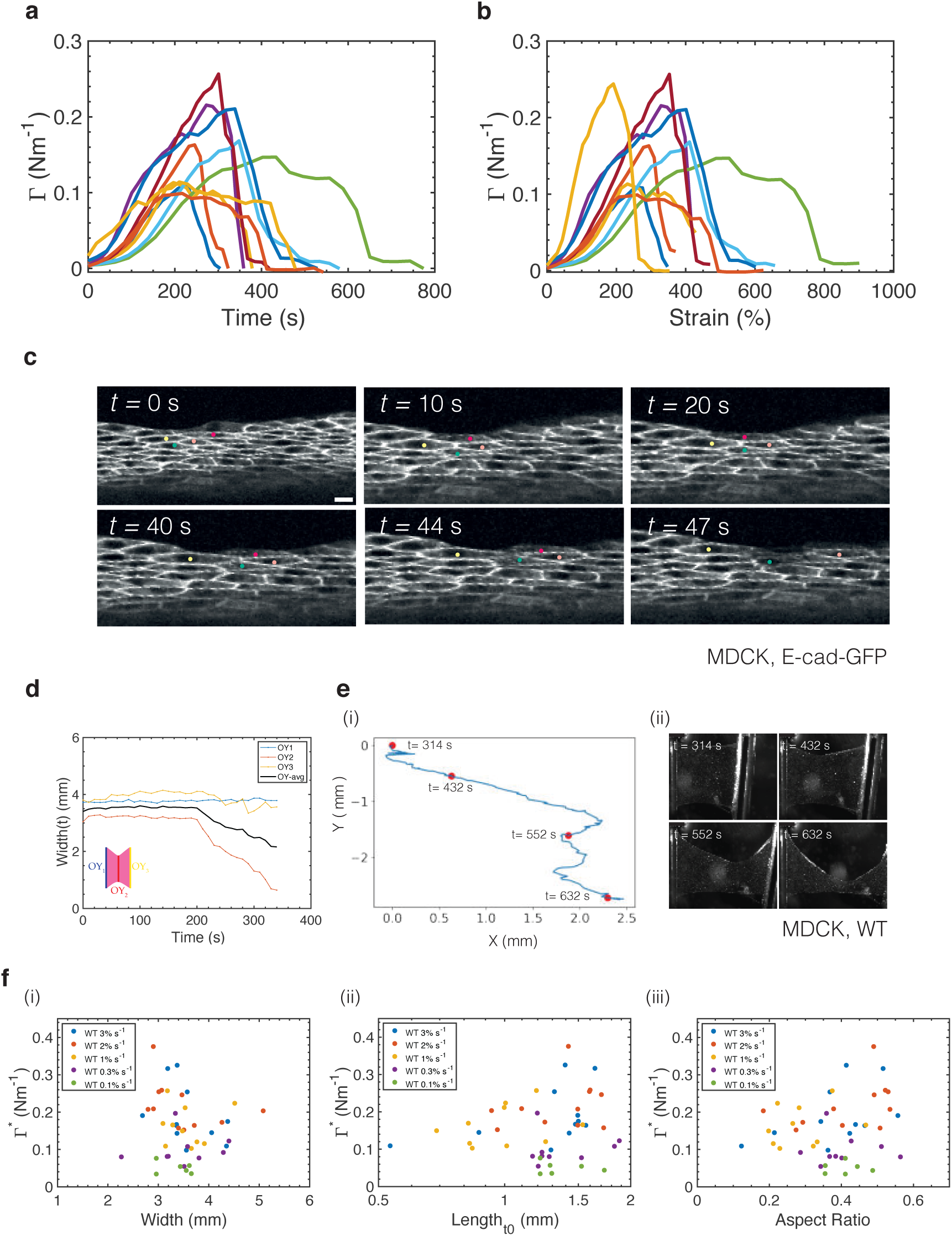
Cells detach from one another during fracture. **(a)** Tension as a function of time and **(b)** Tension as a function of strain for ramp experiments performed at 1% s^−1^ (n = 10 monolayers). Each curve corresponds to a separate monolayer. **(c)** Time series of crack propagation in a monolayer of MDCK cells expressing E-Cadherin-GFP. Several cells are marked by coloured dots in each frame. Time is indicated in the top left corner. Scale bar= 10µm. **(d)** Temporal evolution of monolayer average width for the monolayer shown in **Fig. 1c**. The average width of the monolayer OY_avg_ is shown in black and is computed from the width at each of the arms of the device, OY_1_, OY_3_ and the width at the middle of the monolayer OY_2_. These locations are indicated in the schematic diagram in the bottom le0ft of the graph. The colour of the curves correspond to the position where the width is measured. **(e)** Representative propagation of the crack front in an MDCK monolayer subjected to a strain ramp performed at 1% s^−1^. (i) Crack trajectory in the referential of the camera. Each segment corresponds to 1s. The crack starts at the top left and propagates towards the bottom right. Red dots correspond to images in (ii). The timing of each dot is indicated on the graph. (ii) Time series of crack propagation in an MDCK monolayer. **(f)** Rupture tension Γ as a function of monolayer initial width (i), initial length (ii), and aspect ratio (iii). Each dot corresponds to a separate monolayer. Each colour corresponds to a different strain rate.

**Figure S2:**
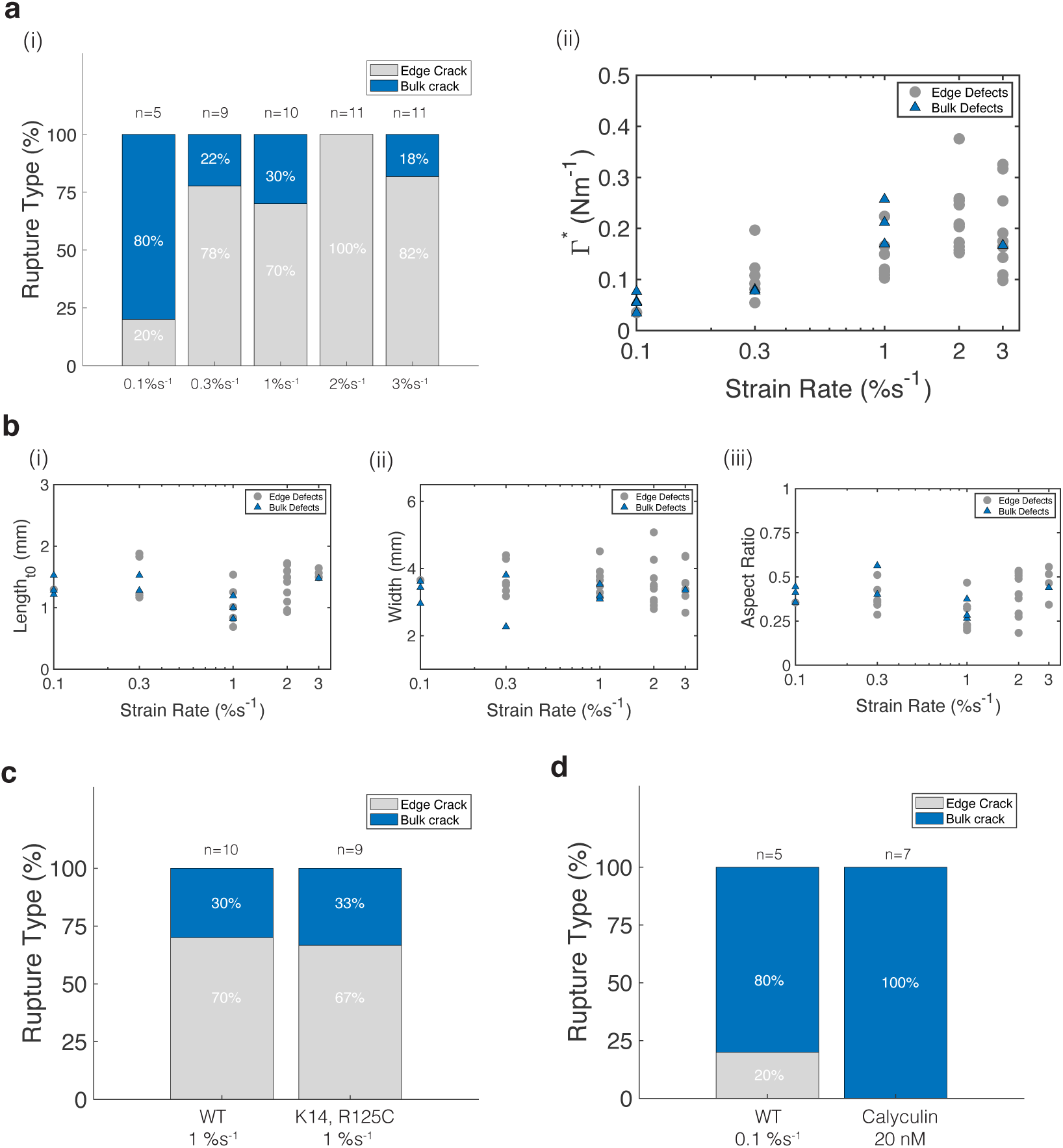
Location of initial crack. In all bar charts, The number of monolayers examined is indicated above each bar. **(a)** Location of initial crack as a function of strain rate. (i) Location of first visible crack (bulk in blue vs edge in grey) as a function of strain rate. (ii) Rupture tension as a function of strain rate. Each marker corresponds to a monolayer and indicates the location of the crack: grey dot for edge defects and black triangles for bulk defects. **(b)** Initial length (i), width (ii) and aspect ratio (iii) as a function of strain rate. Each marker corresponds to a monolayer and indicates the location of the crack: grey dot for edge defects and black triangles for bulk defects. **(c)** Location of the first crack for WT and K14, R125C monolayers stretched at 1% s^−1^ strain rate. **(d)** Location of the first crack for WT monolayers stretched at 0.1% s^−1^ and monolayers treated with 20nM calyculin.

**Figure S3:**
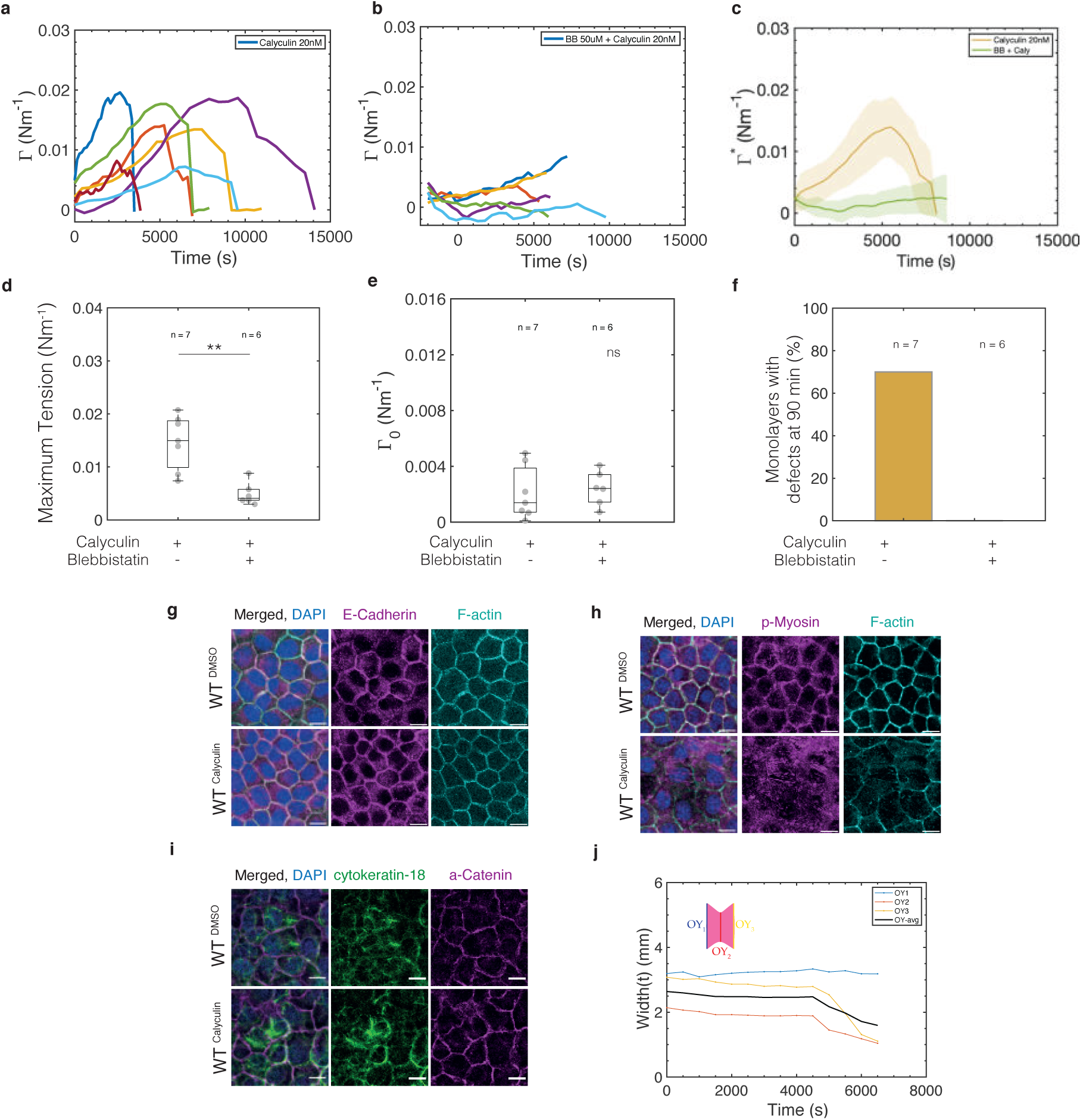
Increase in contractility by calyculin treatment leads to monolayer rupture. In all box plots, the central mark indicates the median, and the bottom and top edges of the box indicate the 25th and 75th percentiles, respectively. The whiskers extend to the most extreme data points that are not outliers. Data points appear as grey dots. Statistically significant difference: ns non significant P *>* 0.05, *P *<* 0.05, ***P *<* 0.001, ****P *<* 0.0001, Kolmogorov-Smirnov test. **(a)** Temporal evolution of tension for calyculin-treated monolayers (n = 7). Each curve corresponds to a separate monolayer. **(b)** Temporal evolution of tension for monolayers treated with blebbistatin and calyculin (n = 6). Each curve corresponds to a separate monolayer. **(c)** Average temporal evolution of tension (thick lines) for monolayers treated with calyculin only (orange, n = 7) or treated with blebbistatin and calyculin (green, n= 6). Tension in monolayers pre-treated with blebbistatin (green) does not increase as in those treated with calyculin (orange). Shaded areas depict the standard deviation. **(d)** Maximum tension reached in monolayers treated with calyculin alone and calyculin+blebbistatin. **(e)** Pre-tension in monolayers in each set of experiments. Pre-tension is measured before adding any drug. **(f)** Percentage of monolayers that show defects during the first 90 min of calyculin treatment for monolayers treated with calyculin alone or calyculin+blebbistatin. (g-i) Immunostainings of wild-type MDCK cells treated with DMSO (first row) or Calyculin 20nM (second row) for 15-20 min. In all panels, the leftmost column shows an overlay. Scale bars, 10 µm. **(g)** Immunostaining against E-Cadherin (magenta) and F-actin (cyan). **(h)** Immunostaining against phospho-Myosin (p-Myosin, magenta) and F-actin (cyan). **(i)** Immunostaining against cytokeratin-18 (green) and alpha-catenin (magenta). **(j)** Temporal evolution of monolayer average width for the monolayer shown in **Fig. 2f**. The average width of the monolayer OY_avg_ is shown in black and is computed from the width at each of the arms of the device, OY_1_, OY_3_ and the width at the middle of the monolayer OY_2_. These locations are indicated in the schematic diagram in the top left of the graph. The colour of the curves correspond to the position where the width is measured.

**Figure S4:**
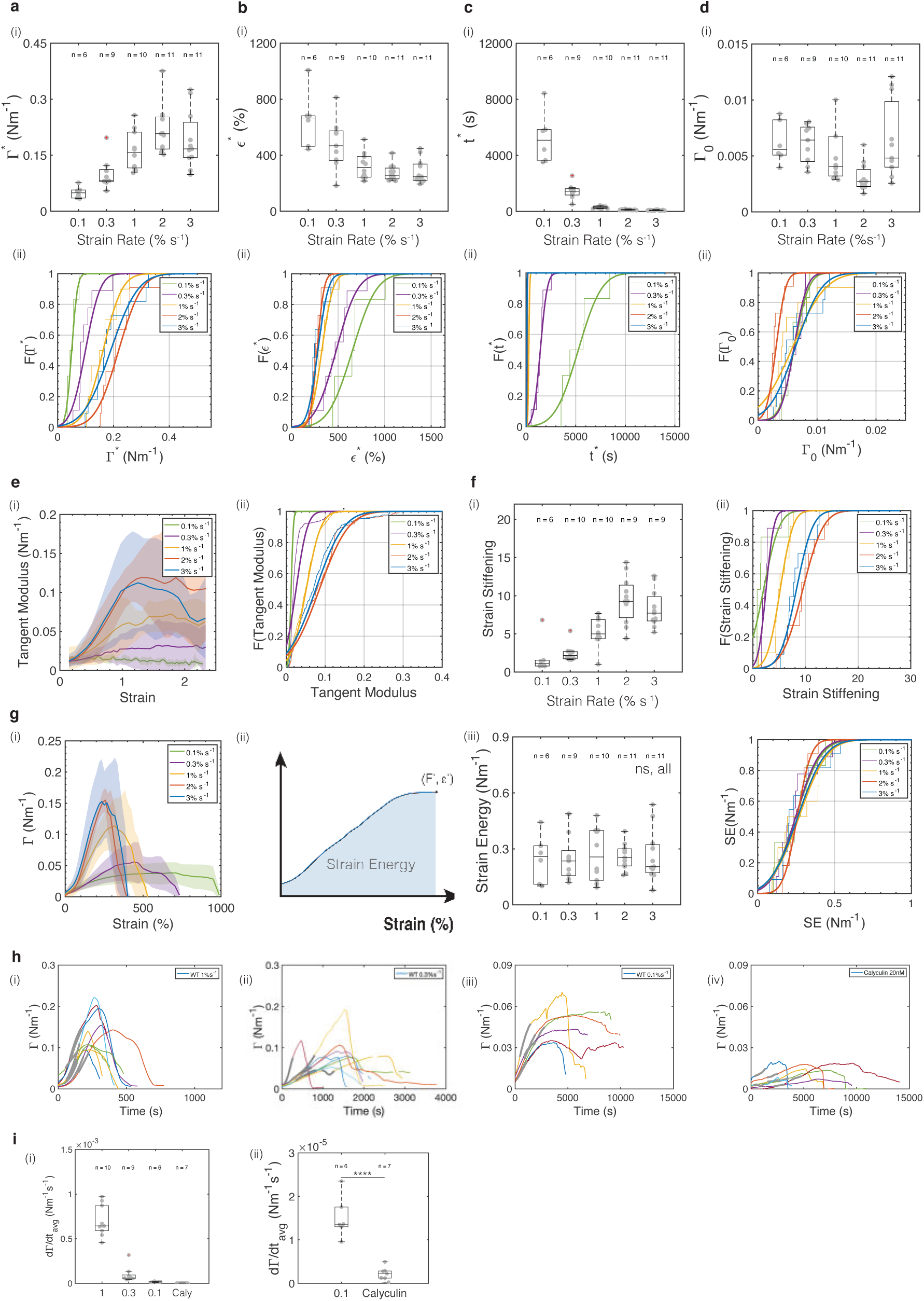
Monolayer rupture characteristics and strain-stiffening scale with strain rate. In all box plots, the central mark indicates the median, and the bottom and top edges of the box indicate the 25th and 75th percentiles, respectively. The whiskers extend to the most extreme data points that are not outliers. Data points appear as grey dots. Outliers are indicated with a red ‘+’ symbol. Statistically significant difference: ns non significant P *>* 0.05, *P *<* 0.05, ***P *<* 0.001, ****P *<*0.0001, Kolmogorov-Smirnov test. Data was acquired from n = 6 monolayers for 0.1% s^−1^, n = 9 for 0.3% s^−1^, n = 10 for 1% s^−1^, n = 11 for 2% s^−1^, and n = 11 for 3% s^−1^. **(a)** Rupture tension, **(b)** rupture strain, **(c)** rupture time and **(d)** pre-tension for different strain rates. **(i)** Box plots. Statistical analysis for these box plots appear in **Tables S1 - S4**. (ii) Cumulative distribution functions *F* (*x*), for the different rupture parameters *x*. Segmented lines represent the empirical cumulative distribution function whereas the solid curves show a fit. Fitting curves were determined using the cumulative distribution function for a Gaussian distribution evaluated on a given segment of the data. The parameters evaluated correspond to the ones shown in panels a-d(i). **(e)** Tangent modulus as a function of strain plotted for all strain rates. **(i)** Graph showing the average value of the gradient (thick lines) and its standard deviation (shaded area) for all strain rates. (ii) Cumulative distribution functions of the tension gradient. **(f)** Fold change in tangent modulus between 15% and 120% strain as a function of strain rate. (i) Box plots of the strain-stiffening between 15% and 120% strain for all strain rates. Statistical analysis appears in **Table S5**. (ii) cumulative distribution functions. **(g)** Strain energy. (i) Evolution of tension as a function of strain for different strain rates. Thick lines indicate the average and shaded areas show the standard deviation. (ii) The strain energy is calculated as the integral of the tension-strain curve up until rupture (Γ^∗^, *ɛ*^∗^). (iii) Left: box plots of strain energy as a function of strain rate. Right: cumulative distribution functions. **(h)** Temporal evolution of tension for wild-type monolayers stretched at (i) 1% s^−1^, (ii) 0.3% s^−1^, (iii) 0.1% s^−1^, and (iv) treated with 20nM calyculin. Each line represents an individual monolayer. (i) Average rate of increase in tension for different strain rates and calyculin. The rate of increase in tension was calculated from the region indicated in grey in each tension-time curve in h. (ii) Same as in (i) but with a scale optimised for 0.1% s^−1^ strain rate and calyculin.

**Figure S5:**
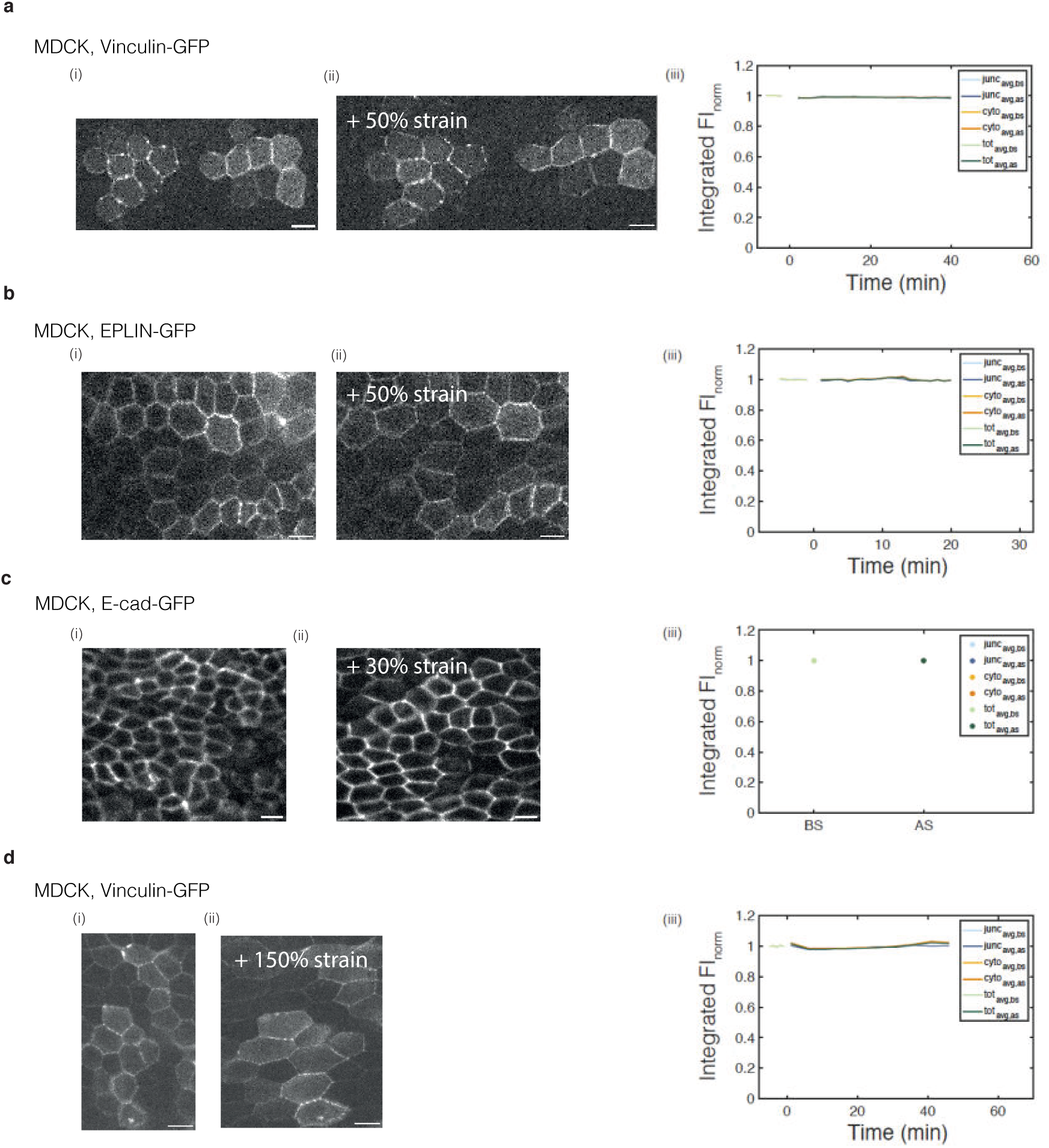
Protein enrichment before and during stretch. Single plane confocal time series of different mechanosensitive proteins before and during stretch. Strain is indicated in the top left corner. Scale bars= 10 µm. **(a)** Vinculin-GFP, (i) before and (ii) after 50% strain applied for 40 min. **(b)** EPLIN-GFP, (i) before and (ii) after 50% strain applied for 20 min. **(c)** E-cadherin-GFP, (i) before and (ii) after 30% strain applied for 40 minutes. **(d)** Vinculin-GFP, (i) before and (ii) after 150% strain applied for 40 min. (a-d) (iii) Temporal evolution of the average fluorescence intensity of the protein of interest. Images were segmented to measure only junctional fluorescence. Stretch is applied at time 0. In the fluorescence intensity graphs, all time points were normalised to the mean value of the average fluorescence intensity before stretch.

**Figure S6:**
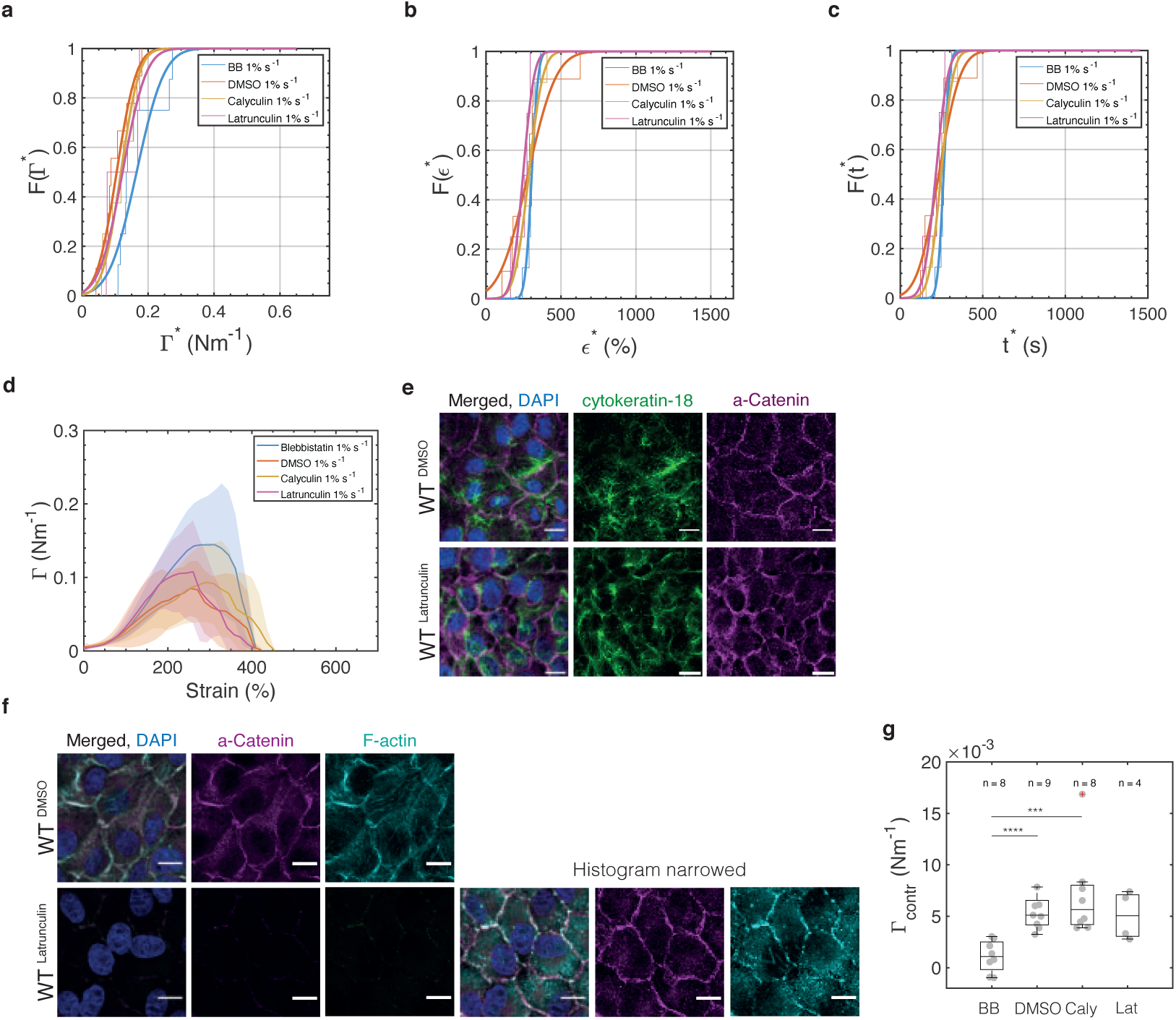
Perturbations of the actin cytoskeleton and myosins do not alter keratins. Cumulative distribution functions for the **(a)** rupture tension, **(b)** rupture strain and **(c)** rupture time in monolayers pre-incubated with different treatments perturbing the actomyosin cytoskeleton. Distributions are computed from the box plots shown in **Fig 4c-e**. **(d)** Tension versus strain curves for monolayers treated with blebbistatin (blue), DMSO (red), calyculin (yellow), and latrunculin (purple). Monolayers were subjected to ramp experiments performed at 1% s^−1^. Thick lines indicate the average and the shading indicates the standard deviation. (a-d, g) Data was acquired from n = 8 monolayers for blebbistatin, n = 9 for DMSO, n = 8 for calyculin, and n = 4 for latrunculin. (e-f) Immunostainings showing the effect of latrunculin 1µM and DMSO. Scale bars, 10 µm. **(e)** Immunostaining against cytokeratin-18 (green) and alpha-catenin (magenta) in WT monolayers treated with DMSO (top row) and latrunculin (bottom row). **(f)** Left: Immunostaining against alpha-Catenin (magenta) and F-actin (cyan) in WT monolayers treated with DMSO (top row) and latrunculin (bottom row). Bottom right: the same images are shown as on the bottom left but with a narrower histogram for alpha-catenin and phalloidin to allow visualisation of the remaining F-actin and alpha-catenin. **(g)** Tension due to contractility in response to different treatments perturbing the actomyosin cytoskeleton. In the box plots, the central mark indicates the median, and the bottom and top edges of the box indicate the 25th and 75th percentiles, respectively. The whiskers extend to the most extreme data points that are not outliers. Data points appear as grey dots. Outliers are indicated with a red ‘+’ symbol. Statistically significant difference: ns non significant: P *>* 0.05, *:P *<* 0.05, ***:P *<* 0.001, ****:P *<* 0.0001, Kolmogorov-Smirnov test.

**Figure S7:**
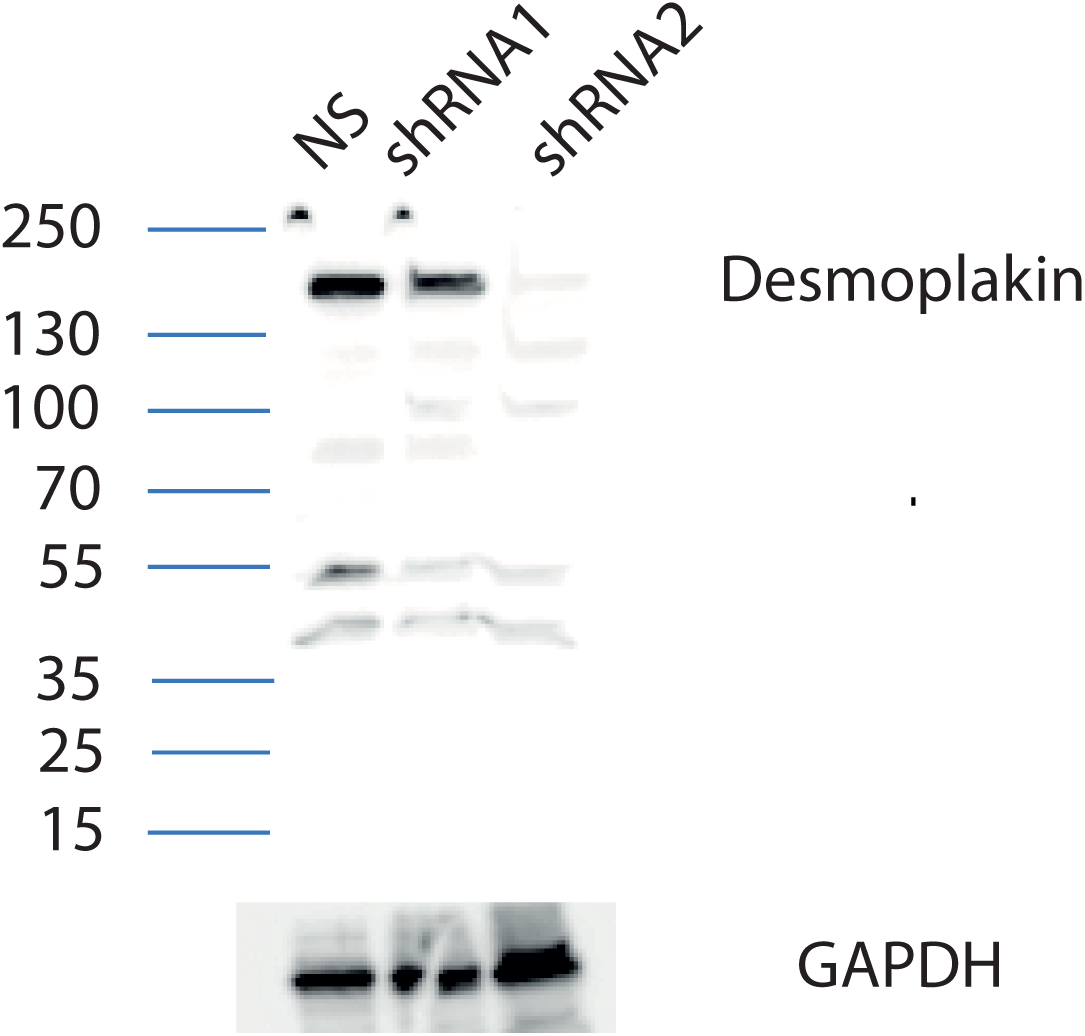
Immunoblot of MDCK cells stably expressing non-silencing (NS) and shRNAs targeting desmoplakin. Reduced levels of desmoplakin expression were observed in MDCK cells expressing shRNA2 compared to cells expressing the non-silencing control plasmid pGIPZ. Immunoblot was probed with anti-desmoplakin and anti-GAPDH antibodies. MDCK cells expressing shRNA2 were used in the experiments presented in **Figs 5**, **6** and **Fig. S8**, **S.9**.

**Figure S8:**
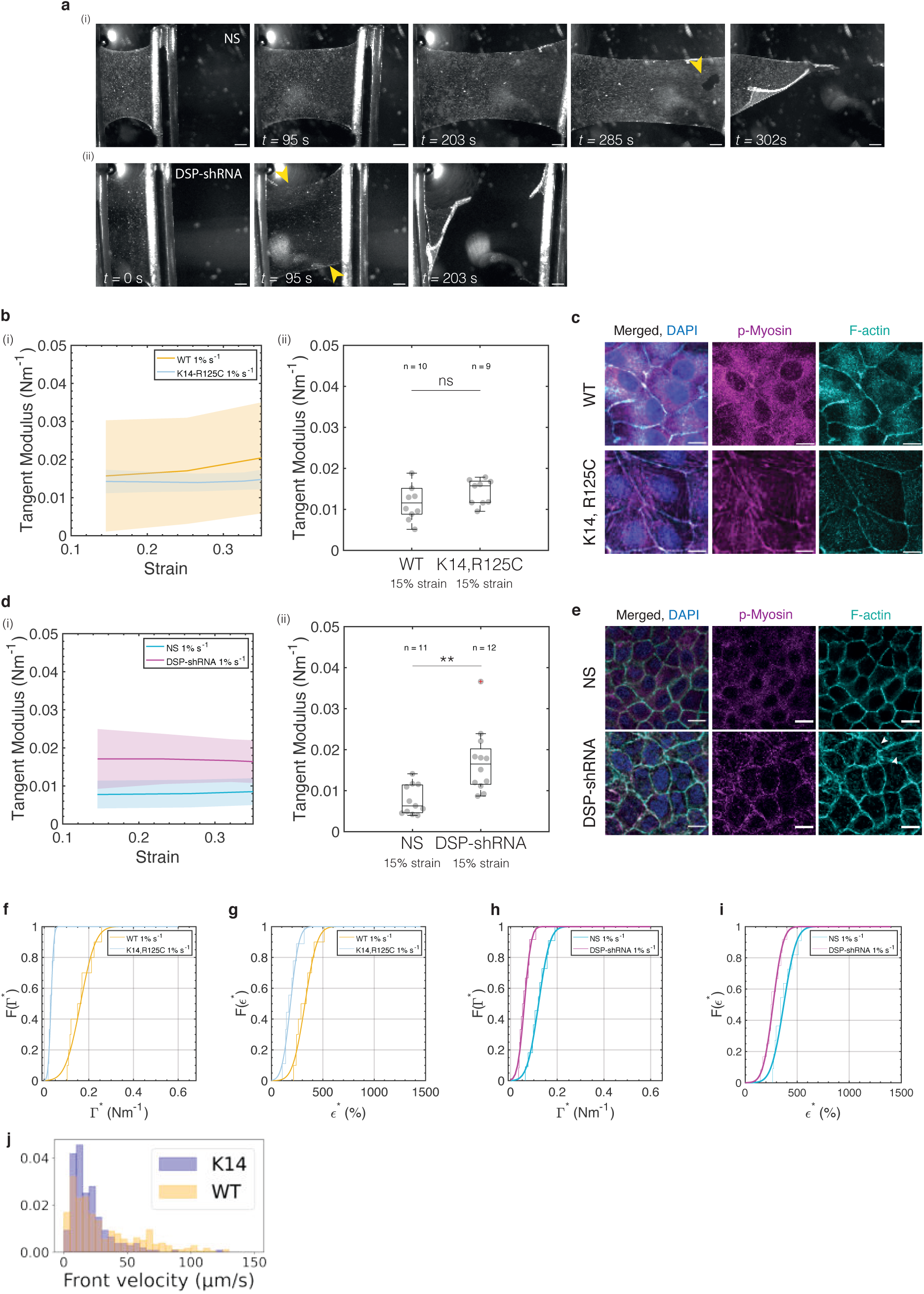
Strain stiffening arises from the intermediate filament cytoskeleton. In all box plots, the central mark indicates the median, and the bottom and top edges of the box indicate the 25th and 75th percentiles, respectively. The whiskers extend to the most extreme data points that are not outliers. Data points appear as grey dots. Outliers are indicated with a red ‘+’ symbol. Statistically significant difference: ns non significant: P *>* 0.05, *:P *<* 0.05, ***:P *<* 0.001, ****:P *<* 0.0001, Kolmogorov-Smirnov test. **(a)** Bright-field time series of representative **(i)** non-silencing shRNA (NS) and (ii) desmoplakin-shRNA (DSP-shRNA) monolayers during a ramp experiment performed at 1% s^−1^. Arrowheads indicate the onset of rupture. **(b)** Tangent modulus at low strain for WT (yellow, n = 10) and K14, R125C (blue, n = 9) monolayers subjected to a ramp at 1% s^−1^. **(i)** Tangent modulus as a function of strain. Solid lines indicate the average value of the tension gradient over strain and shaded areas show the standard deviation. (ii) Box plots of the tangent modulus at 15% strain in WT and K14, R125C monolayers. The number of monolayers examined is indicated above the box plots. **(c)** Immunostaining against p-myosin (magenta) and F-actin (cyan) for WT (top row) and K14, R125C (bottom row) monolayers. Scale bar= 10µm. **(d)** Tangent modulus at low strain in NS control (cyan, n= 11) and DSP-shRNA (magenta, n = 12) monolayers subjected to a ramp in deformation at 1% s^−1^. Tangent modulus as a function of strain. Solid lines represent the average value and shaded areas show the standard deviation of the distribution. (ii) Box plots of the tangent modulus at 15% strain for NS control and DSP-shRNA monolayers. **(e)** Immunostaining against p-myosin (magenta) and F-actin (cyan) for NS-shRNA (top row) and DSP-shRNA (bottom row) monolayers. Scale bar= 10µm. (f-i) Cumulative distribution functions computed from the box plots in **Fig. 5e-h**. (f-g) Cumulative distribution function for the rupture tension **(f)** and rupture strain **(g)** for WT and K14, R125C monolayers. (h-i) Cumulative distribution functions of the rupture tension **(h)** and rupture strain **(i)** for NS-shRNA control and DSP-shRNA monolayers. **(j)** Distribution of crack front velocities for wild-type monolayers (WT, orange) and K14, R125C monolayers (K14, purple).

**Figure S9:**
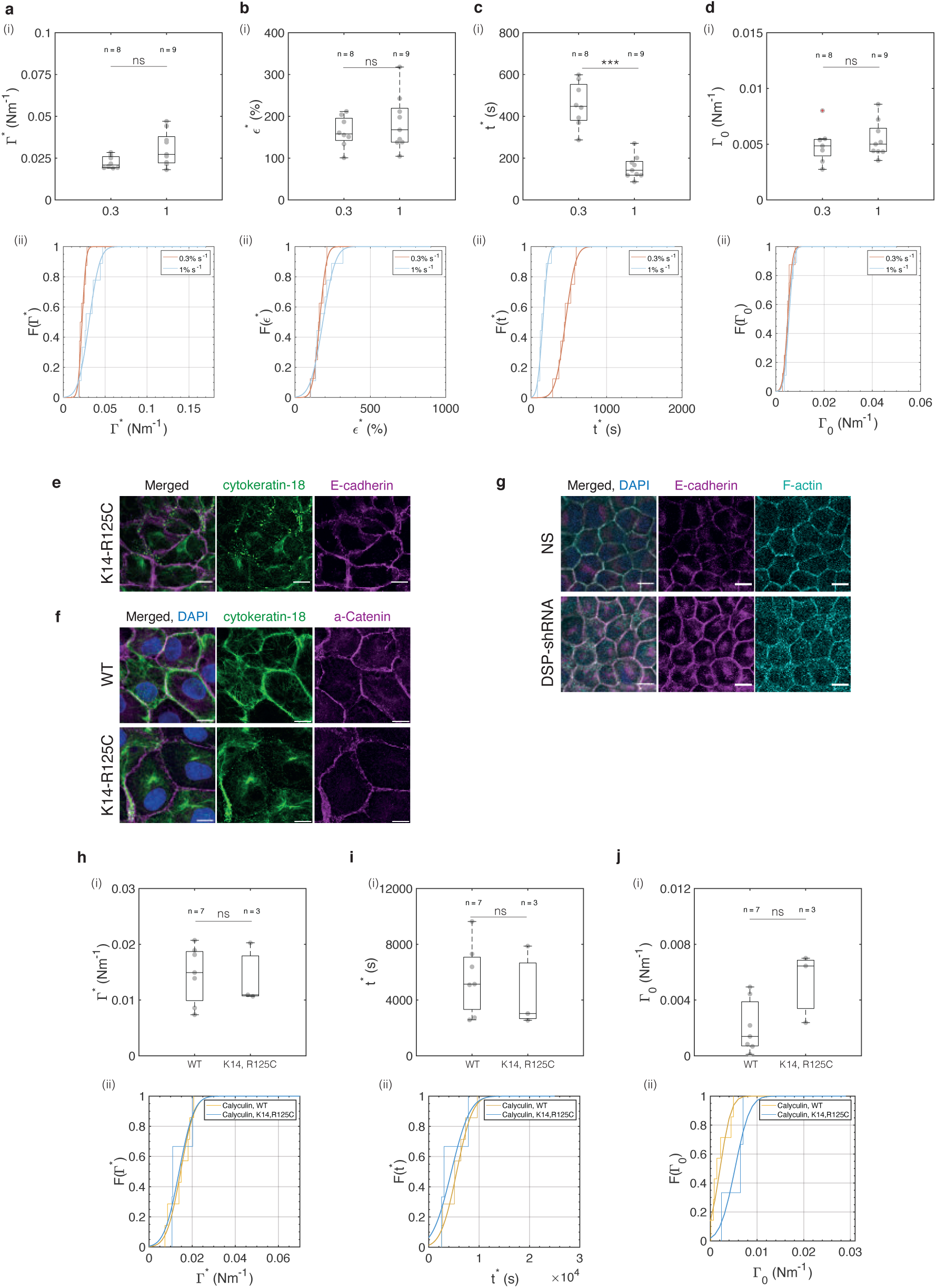
Rupture characteristics do not depend on strain rate when keratin intermediate filaments are perturbed. In all box plots, the central mark indicates the median, and the bottom and top edges of the box indicate the 25th and 75th percentiles, respectively. The whiskers extend to the most extreme data points that are not outliers. Data points appear as grey dots. Outliers are indicated with a red ‘+’ symbol. Statistically significant difference, ns non significant: P *>* 0.05, *:P *<* 0.05, ***:P *<* 0.001, ****:P *<* 0.0001, Kolmogorov-Smirnov test. **(a)** Rupture tension, **(b)** rupture strain, **(c)** rupture time, and **(d)** pre-tension in K14, R125C monolayers subjected to a ramp in deformation at 0.3% s^−1^ and 1% s^−1^. (i) Box plots. The number of monolayers is indicated above each box. Cumulative distribution functions. In these plots, data from monolayers subjected to ramps with a strain rate of 0.3% s^−1^ are shown in orange and those at 1% s^−1^ are shown in blue. (e-g) Left panels are merges of middle and right panels. In some, DAPI nuclear staining is also overlaid. Scale bar= 10µm. **(e)** Immunostaining of cytokeratin-18 (green) and E-cadherin (magenta) in K14, R125C monolayers. **(f)** Immunostaining of cytokeratin-18 (green) and alpha-Catenin (magenta) in wild-type (WT, top row) and K14-R125C (bottom row) monolayers. **(g)** Immunostaining of E-cadherin (magenta) and F-actin (cyan) in NS control (top row) and DSP-shRNA monolayers (bottom row). **(h)** Rupture tension, **(i)** rupture time, and **(j)** pre-tension in WT and K14, R125C monolayers treated with 20nM calyculin. **(i)** Box plots. The number of monolayers is indicated above each box. (ii) Cumulative distribution functions computed from box plots in (i). In these plots, data from WT monolayers treated with calyculin are shown in orange and those from K14, R125C monolayers treated with calyculin are shown in blue.

**Figure S10:**
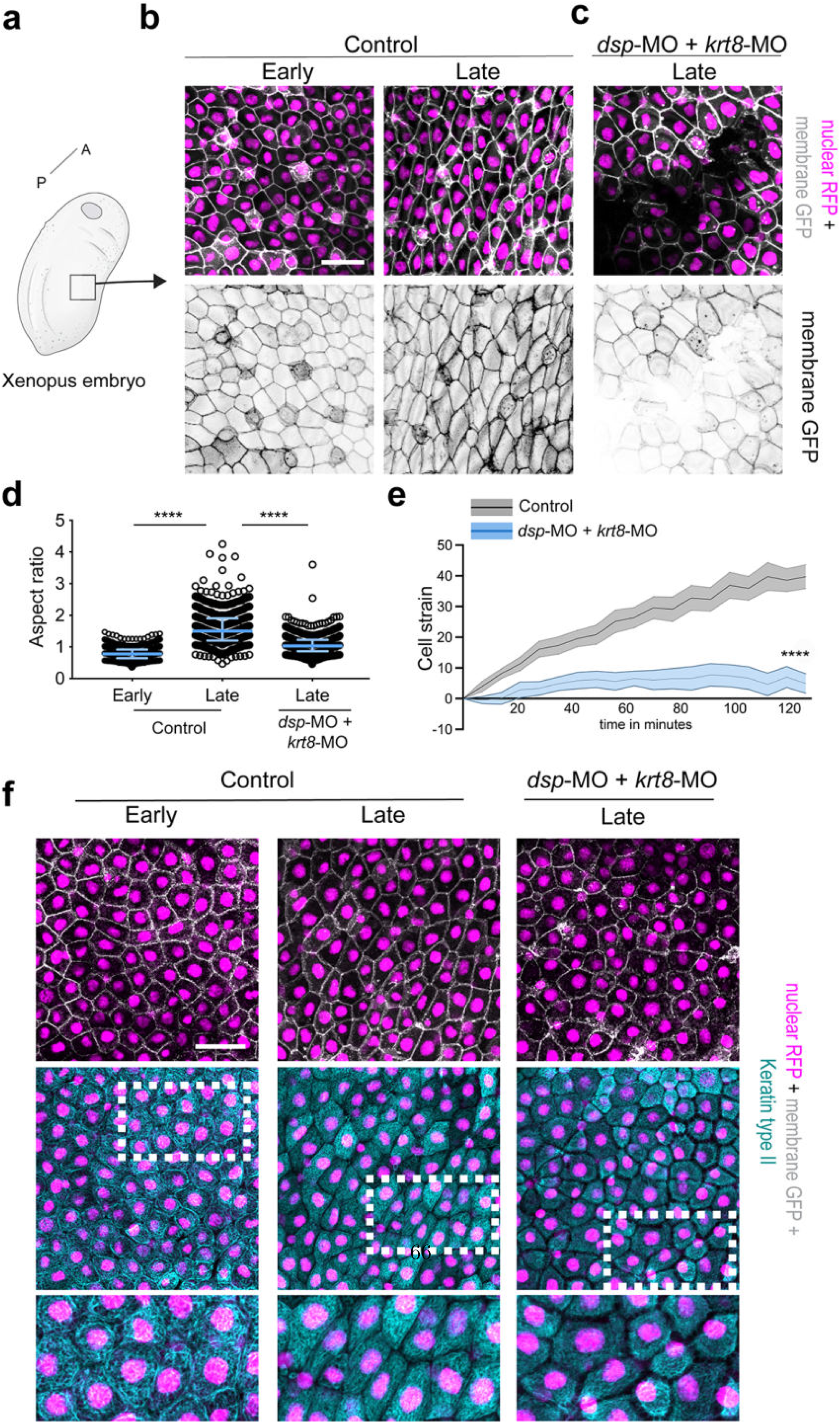
Perturbation of the keratin intermediate filament network in *Xenopus laevis* embryos leads to tissue rupture during development. **(a)** Schematic representation of a *Xenopus laevis* embryo. A, anterior; P, posterior. The black box in the lateral epidermis indicates the region of interest imaged in b, c, and f. **(b)** Representative confocal projections of the epidermal cells at early (stage 16) and late (stage 21-22) stages of body axis elongation. **(c)** Representative confocal projections showing the impact of desmoplakin and keratin 8 knockdown in the lateral epidermis at late stages of body axis elongation. (b, c) Top row: composite image showing the nucleus in magenta and the membrane in white. Bottom row: membrane GFP. **(d)** Aspect ratio in early and late control embryos and in late embryos depleted in desmoplakin and keratin 8. The bar represents the median and the whiskers indicate the interquartile ranges; Kruskal-Wallis test ****:P *<* 0.0001, individual comparisons Mann Whitney test ****P *<* 0.0001. **(e)** Cell strain as a function of time for control (black) and knockdown (blue) embryos. Solid lines represent the average value and shaded areas show the standard deviation of the distribution. Unpaired two-tailed t-test **** P*<* 0.0001. **(f)** Confocal projections of embryos immunostained against keratin type II. Left: control embryos at early and late stages. Right: Embryos depleted in desmoplakin and keratin 8 at late stages of body axis elongation. Top row: overlay of membrane-GFP (white) and nucleic acids (magenta). Middle row: Overlay of cytokeratin (cyan, immunostaining) and nucleic acids (magenta). Bottom row: Zoom of the region in the dashed white box in the middle row. Scale bars in b, c, and f: 40 µm. b,c,f are representative examples of at least three independent experiments, C.I. 95%.

**Figure S11:**
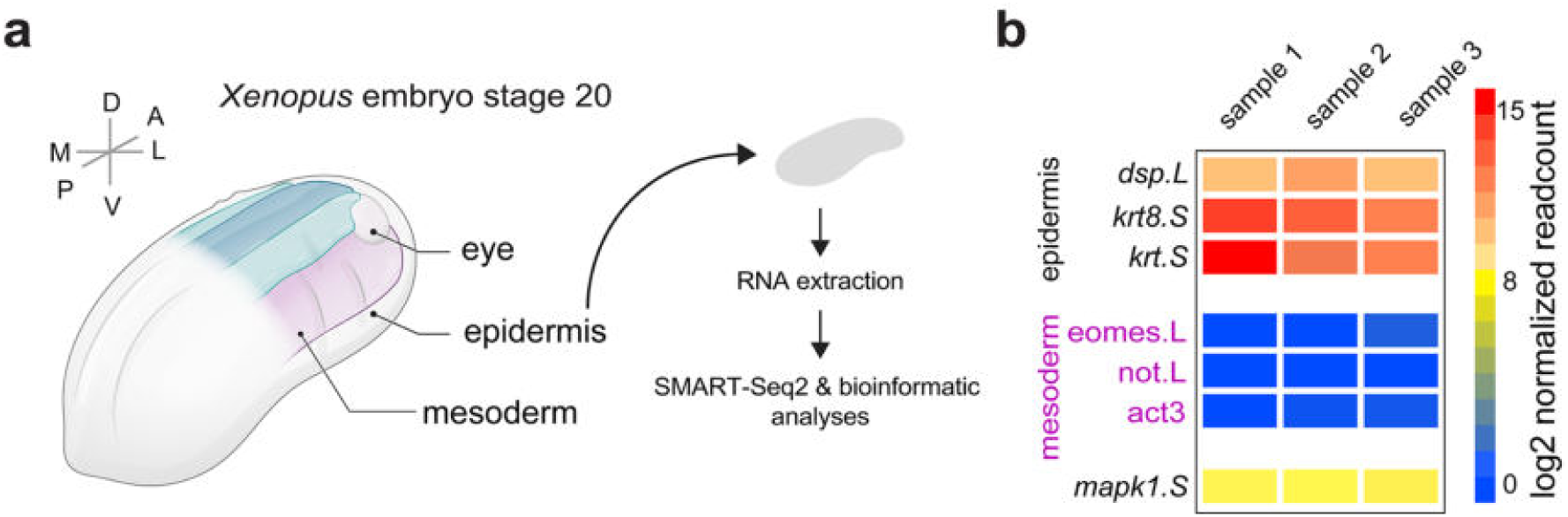
Transcriptomics shows that desmoplakin and keratin 8 are highly expressd in *Xenopus laevis* epidermis. **(a)** Schematic representation of the anatomy of a *Xenopus laevis* embryo. Epidermis was collected from embryos and processed for RNA extraction, SMART-Seq2 and bioinformatics analysis. **(b)** Sample quality control showing that desmoplakin (dsp.L0 and keratin 8 (krt8.S) are enriched in libraries from all epidermis samples. Mesoderm-specific genes (magenta) are present at low levels confirming that most of the cells are epidermal. The final line (mapk1.S) shows a housekeeping gene. Three independent experiments are shown.

**Figure S12:**
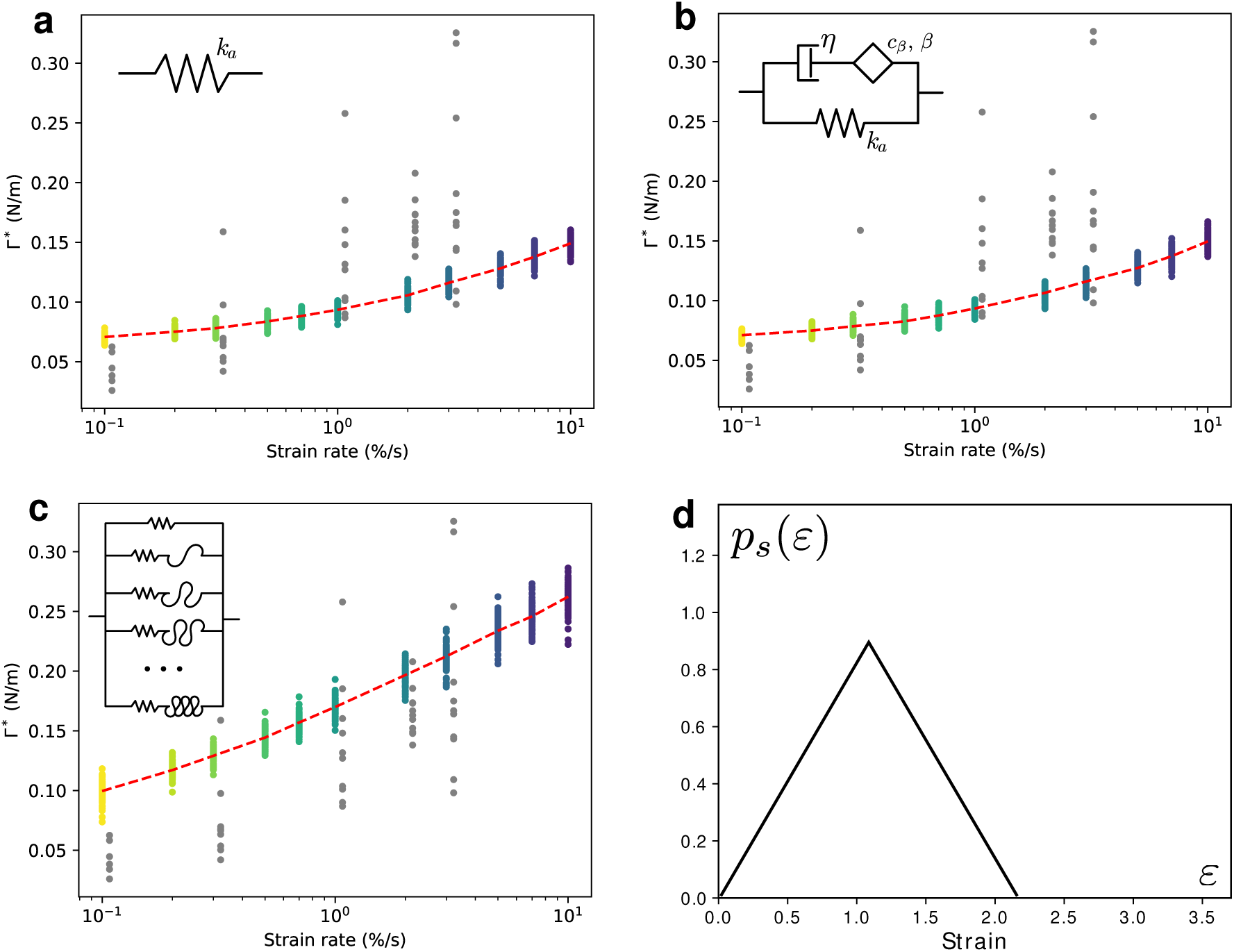
Multiscale modelling of fracture onset. Each coloured dot represents a simulation run. 100 simulations were run for each strain rate. Each grey dot represents an experimental data point. The dashed and dotted red lines link the mean value for each strain rate to show the trend. **a - c**, Rupture tension as a function of the strain rate for the three rheological models presented in **Fig. 7 d, e, g** respectively. **d**, Distribution *p_s_* of strains at which keratin bundles are recruited to bear load.

**Figure S13:**
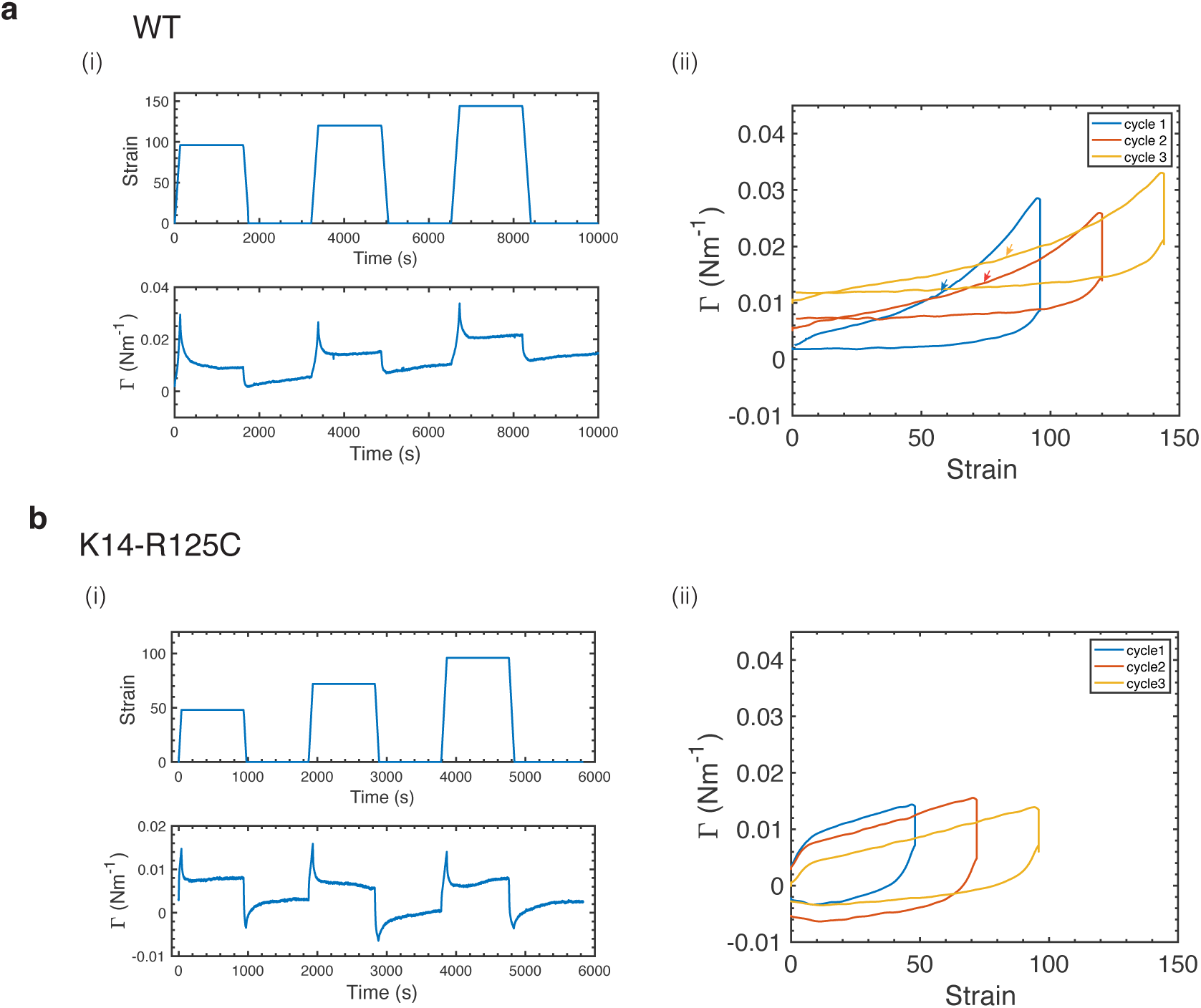
The strain stiffening threshold depends on strain history. **(a)** Wild type monolayer subjected to successive cycles of stretch and unloading at 1% s^−1^ strain rate. **(i)** Top: Strain imposed as a function of time. Bottom: Tension as a function of time. (ii) Tension as a function of strain for all cycles. Each cycle appears in a different colour and an arrow indicates the threshold of strain stiffening. **(b)** K14-R125C monolayers were subjected to successive cycles of stretch and unloading at 1% s^−1^ strain rate. (i) Top: Strain imposed as a function of time. Bottom: Tension as a function of time. (ii) Tension as a function of strain for all cycles. Each cycle appears in a different colour.

**Figure S14:**
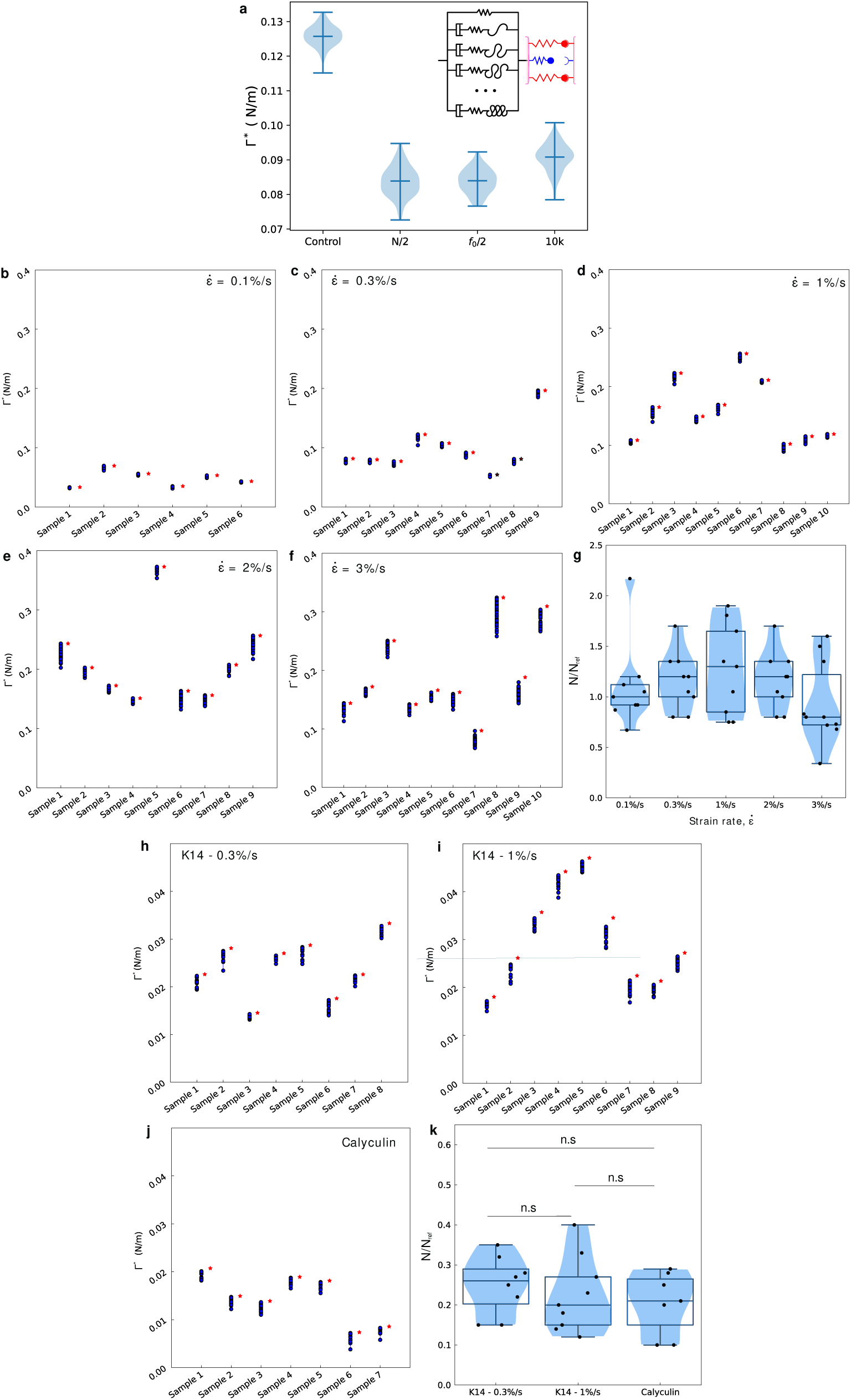
Relative linker number statistics for WT, K14 mutant and calyculin treated tissues. **(a)** Violin plots showing the effect of varying the bond model parameters on the resulting rupture tension Γ^∗^, for an imposed ramp at 1% s^−1^. The middle line shows the median and the whiskers the extremes. The outcome of the WT model parameters are compared to the outcome of the following situations: reducing the number of bonds per-junction *N* by a factor 2 (N/2), reducing the force threshold *f*_0_ in the slip bond behaviour by a factor two (*f*_0_*/*2), or increasing the rates *k_on_* and *k_off,_*_0_ by a factor (10k) all lead to a significant reduction of the rupture tension. (b-f) Distributions of rupture tensions for the bond model when fitting the WT stress time series, experiment by experiment, by adjusting the linker number *N* while keeping all the other model parameters at their reference values (see **Table S7**). The red asterisks show individual experimental measurements, and blue dots show simulated rupture points for the optimal N value. **(g)** Summary statistics of the resulting *N* distributions across all loading rates in WT monolayers. Each black dot represents the analysis of an individual monolayer. (h-j) Similar to **(a)** but now for monolayers expressing K14-R125C (h-i) and calyculin-treated WT monolayers (j). **(k)** Sumary distributions of *N* values for K14-R125C monolayers and calyculin-treated WT monolayers. Each black dot represents the analysis of an individual monolayer.

## Statistical Analysis for ramps performed at different strain rates in wild-type monolayers

**Table S1:**
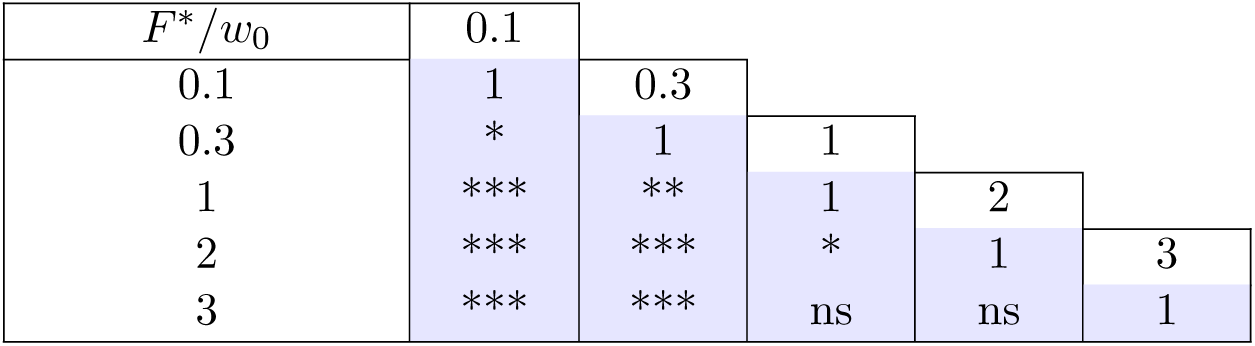
Statistical analysis of the rupture tension of monolayers subjected to ramps in deformation at different strain rates (0.1 % s^−1^ - 3 % s^−1^). Statistically significant difference was determined using a Kolmogorov-Smirnov test: ns P *>* 0.05, *P *<* 0.05, **P *<*0.01, ***P *<* 0.001.

| $F^*/w_0$ | 0.1 | | | | | |
| --- | --- | --- | --- | --- | --- | --- |
| 0.1 | 1 | 0.3 |  |  |  |  |
| 0.3 | * | 1 | 1 |  |  |  |
| 1 | *** | ** | 1 | 2 |  |  |
| 2 | *** | *** | * | 1 | 3 |  |
| 3 | *** | *** | ns | ns | 1 |  |

**Table S2:**
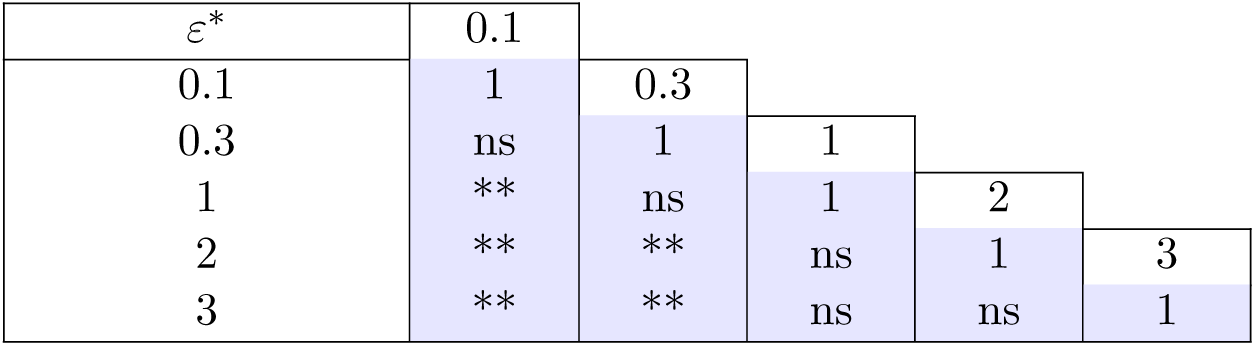
Statistical analysis of the rupture strain of monolayers subjected to ramps in deformation at different strain rates (0.1 % s^−1^ - 3 % s^−1^). Statistically significant difference was determined using a Kolmogorov-Smirnov test: ns P *>* 0.05, **P *<* 0.01.

| $\varepsilon^*$ | 0.1 | | | | | |
| --- | --- | --- | --- | --- | --- | --- |
| 0.1 | 1 | 0.3 |  |  |  |  |
| 0.3 | ns | 1 | 1 |  |  |  |
| 1 | ** | ns | 1 | 2 |  |  |
| 2 | ** | ** | ns | 1 | 3 |  |
| 3 | ** | ** | ns | ns | 1 |  |

**Table S3:** Statistical analysis of the rupture time of monolayers subjected to ramps in deformation at different strain rates (0.1 % s^−1^ - 3 % s^−1^). Statistically significant difference was determined using a Kolmogorov-Smirnov test: *P *<* 0.05, ***P *<* 0.001, ****P *<* 0.0001.

| $t^*$ | 0.1 | | | | | |
| --- | --- | --- | --- | --- | --- | --- |
| 0.1 | 1 | 0.3 |  |  |  |  |
| 0.3 | *** | 1 | 1 |  |  |  |
| 1 | *** | **** | 1 | 2 |  |  |
| 2 | *** | **** | *** | 1 | 3 |  |
| 3 | *** | **** | **** | * | 1 |  |

**Table S4:** Statistical analysis of the pre-tension of monolayers subjected to ramps in deformation at different strain rates (0.1 % s^−1^ - 3 % s^−1^). Statistically significant difference was determined using a Kolmogorov-Smirnov test: ns P *>* 0.05, *P *<* 0.05.

| $F_0/w_0$ | 0.1 | | | | |
| --- | --- | --- | --- | --- | --- |
| 0.1 | 1 | 0.3 |  |  |  |
| 0.3 | ns | 1 | 1 |  |  |
| 1 | ns | ns | 1 | 2 |  |
| 2 | * | * | * | 1 | 3 |
| 3 | ns | ns | ns | ns | 1 |

**Table S5:**
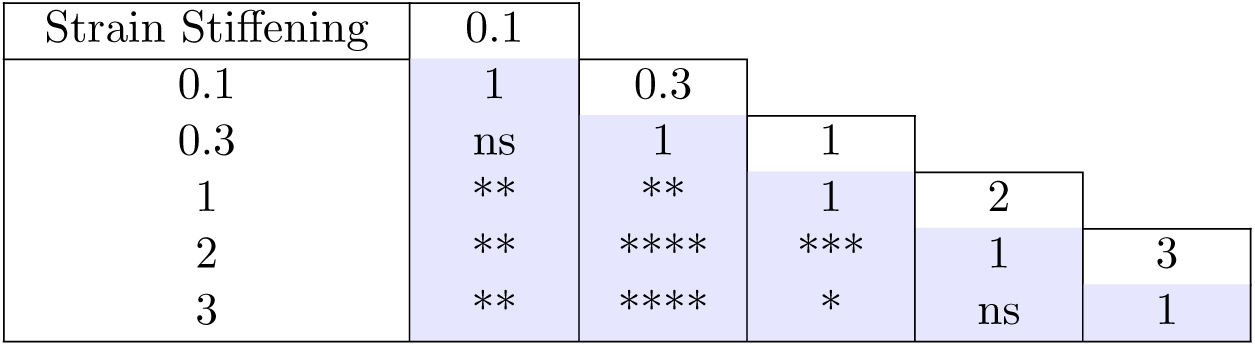
Statistical analysis of the strain stiffening between 15% strain and 120% strain displayed by monolayers subjected to ramps in deformation at different strain rates (0.1 % s^−1^ - 3 % s^−1^). Statistically significant difference was determined using a Kolmogorov-Smirnov test: ns P *>* 0.05, *P *<* 0.05, **P *<* 0.01, ***P *<* 0.001, ****P *<* 0.0001.

| Strain Stiffening | 0.1 |  |  |  |  |
| --- | --- | --- | --- | --- | --- |
| 0.1 | 1 | 0.3 |  |  |  |
| 0.3 | ns | 1 | 1 |  |  |
| 1 | ** | ** | 1 | 2 |  |
| 2 | ** | **** | *** | 1 | 3 |
| 3 | ** | **** | * | ns | 1 |

**Table S6:**
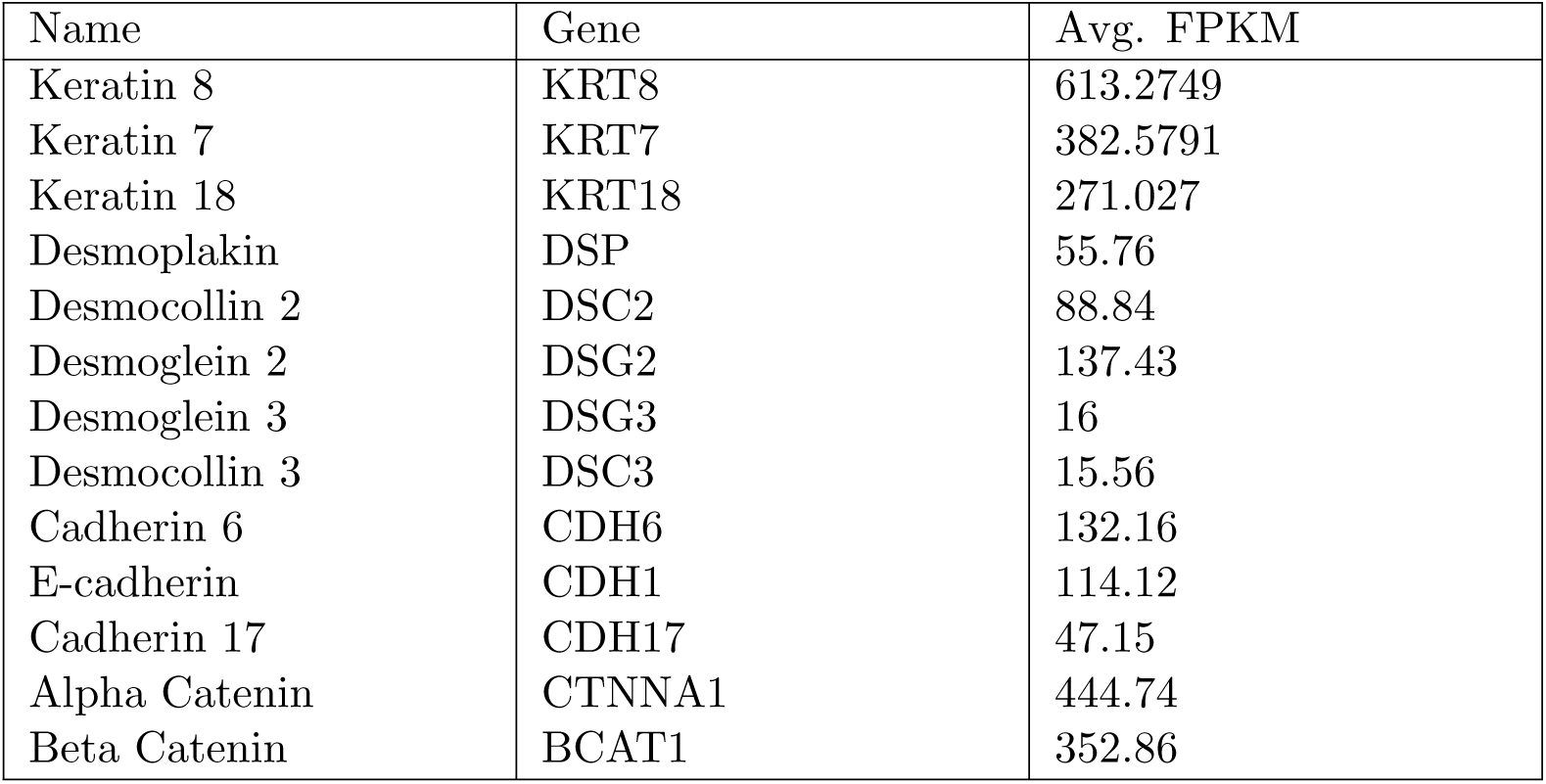
RNA-seq analysis in MDCK cells. The first column indicates the names of the proteins, in the second column appear the gene names, and the third column shows the mRNA abundance obtained by RNA-seq in fragments per kilobase million (FPKM).

**Table S7:** Parameters for the binding-unbinding model.

|  |  |
| --- | --- |
| Number of bonds, $N_{\text{ref}}$ | 100 |
| $k_{on}$ (1/s) | 3e-3 |
| $k_{off,0}$ (1/s) | 3e-4 |
| $f_0$ (N) | 0.055 |

**Table S8:** Parameters for the four material models shown in **Fig. 7**.

| Linear elastic model |  |  |  |
| --- | --- | --- | --- |
| $k_a$ (Pa) | - | - | - |
| 760 | - | - | - |
| Linear viscoelastic model |  |  |  |
| $k_a$ (Pa) | $c_\beta$ (Pa · s$^\beta$) | $\beta$ | $\eta$ (Pa · s) |
| 760 | 1360 | 0.22 | 1e4 |
| Nonlinear spring |  |  |  |
| $k$ (Pa) | $\epsilon_{start}$ | $\epsilon_{end}$ | $k_a$ (Pa) |
| 15e3 | 0 | 2.2 | 1e3 |
| Nonlinear viscoelastic model |  |  |  |
| $\tau$ (s) | $\epsilon_{start}$ | $\epsilon_{end}$ | $k_a$ (Pa) |
| 55 | 0 | 2.2 | 1e3 |

## Supplementary Movies

**Supplementary Video 1.** Suspended MDCK wild-type monolayer subjected to a ramp in deformation at 1% s^−1^. Time 0 corresponds to the onset of stretch. Time is in seconds. Scale bar is 500 µm.

**Supplementary Video 2.** Suspended MDCK E-cad-GFP monolayer subjected to constant stretch imaged by phase contrast microscopy at 40X high magnification. Time is in seconds. Scale bar is 10 µm.

**Supplementary Video 3.** Suspended MDCK wild-type monolayer treated with calyculin 20 nM. Calyculin is added at time 0. Time is in seconds. Scale bar is 500 µm.

**Supplementary Video 4.** Suspended MDCK wild-type monolayer pretreated with blebbistatin 50 µM for 30 min. Blebbistatin is added at time 0. Time is in seconds. Scale bar is 500 µm.

**Supplementary Video 5.** Suspended MDCK wild-type monolayer treated with calyculin. The monolayer was previously incubated with blebbistatin for 30 min as shown in Supplementary Video 4. Calyculin is added at time 0. Time is in seconds. Scale bar is 500 µm.

**Supplementary Video 6.** Suspended MDCK wild-type monolayer treated with blebbistatin 50 µM for 20 min and subjected to a ramp in deformation at 1% s^−1^ starting at time 0. Time is in seconds. Scale bar is 500 µm.

**Supplementary Video 7.** Suspended MDCK wild-type monolayer treated with DMSO for 20 min and subjected to a ramp in deformation at 1% s^−1^ starting at time 0. Time is in seconds. Scale bar is 500 µm.

**Supplementary Video 8.** Suspended MDCK wild-type monolayer treated with calyculin 20 nM for 20 min and subjected to a ramp in deformation at 1% s^−1^ starting at time 0. Drug added at time 0. Time is in seconds. Scale bar is 500 µm.

**Supplementary Video 9.** Suspended MDCK wild-type monolayer treated with Latrunculin A 1µM for 20 min and subjected to a ramp in deformation at 1% s^−1^ starting at time 0. Time is in seconds. Scale bar is 500 µm.

**Supplementary Video 10.** Suspended MDCK monolayer overexpressing Keratin 14-R125C subjected to a ramp in deformation at 1% s^−1^. Time 0 corresponds to the onset of stretch. Time is in seconds. Scale bar is 500 µm.

**Supplementary Video 11.** Suspended MDCK non-silencing shRNA monolayer subjected to a ramp in deformation at 1% s^−1^. Time 0 corresponds to the onset of stretch. Time is in seconds. Scale bar is 500 µm.

**Supplementary Video 12.** Suspended MDCK Desmoplakin-shRNA monolayer subjected to a ramp in deformation at 1% s^−1^. Time 0 corresponds to the onset of stretch. Time is in seconds. Scale bar is 500 µm.

**Supplementary Video 13.** Body axis elongation in a *Xenopus laevis* embryo. The anteroposterior (AP) axis is oriented vertically. Lateral epidermis cells expressing a GFP membrane marker (black) elongate as development proceeds, see right hand side of the embryo. Frame interval 7.5 minutes and 20 frames are shown. Scale bar is 500 µm. Representative example of three independent experiments.

**Supplementary Video 14.** Effect of the impact of desmoplakin and keratin 8 knockdown on cell shape during during body axis elongation. Cells express a GFP membrane marker (black). Left: cells within control embryos. Middle: cells within an embryo microinjected with morpholinos against desmoplakin and keratin 8 displaying a mild phenotype. Right: cells with an embryo microinjected with morpholinos against desmoplakin and keratin 8 displaying a severe phenotype. Frame interval 7.5 minutes and 20 frames are shown. Scale bar is 50 µm. Representative of three independent experiments.

